## Supplementary Information and Figures for "Ancient DNA reveals diverse community organizations in the 5th millennium BCE Carpathian Basin"

### Index

### Supplementary Note 1: Description of the archaeological sites

*Aszód-Papi földek*

Aszód-Papi földek is a Late Neolithic (LN) site in the Gödöllő hillside, Northern Hungary. Nándor Kalicz excavated the site between 1960 and 1994 when 0.5 ha of the site was revealed. Among the settlement features 224 graves were found. Based on the excavation, field survey and disturbances caused by the construction of modern buildings, the total size of the site is estimated to be 35-40 ha[1]. The burials formed grave-groups and can be found throughout the entire territory of the settlement. The settlement can be dated from 4800-4730 cal BCE (68.3%) to 4760-4690 cal BCE (68.3%) and the burial activity from 4760-4700 cal BCE (68.3%) to 4710-4640 cal BCE (68.3%) based on Bayesian modeled AMS radiocarbon data (Supplementary Fig. 21, Supplementary Data 2).

The material culture reflects intensive interaction with both the Great Hungarian Plain (GHP) and Transdanubia. Culturally, the Aszód-Papi földek site belongs to the East Transdanubian group of the Lengyel complex but both Lengyel- and Tisza-style pottery appear in the site[1–5]. In the determinable cases children were buried in 42.1% of the graves, females in 35.4% and males in 22.5%[1,6] (Supplementary Fig. 12A). The overwhelming majority of the graves was inhumation, however cremations and symbolic burials also occurred in some cases. The orientation of the burials was varied and the majority of the deceased was laid on their right side. Grave goods expressed age, gender and rank differences. The variety of grave goods was significant, pottery, stone and bone tools and ornaments occurred frequently. Polished stone mace heads, wild boar tusk pendants, and wild boar mandibles were male status goods expressing the role of leading position. *Spondylus* and stone ornaments were prestige goods[1].

*Budapest-Albertfalva-Hunyadi János út*

The site is situated in the western bank of the Danube River in southern Budapest. In 2001-2002, during the excavation of an Early Bronze Age Bell Beaker settlement, some Early Copper Age (ECA) features were also unearthed, which were mainly concentrated in the Northwestern part of the area[7]. Of the four features containing human remains, two were crouched burials lying on their left side (Features 345 and 1037), and two (Features 634 and 1088) were round pits, which, in addition to fragmentary human skeletal remains, contained animal bones, Ludanice-style sherds, stone tools and a grinding stone[7,8] (Supplementary Data 2). Based on the Bayesian-modeled AMS dating of the four human remains, the burials can be dated from 4395-4180 cal BCE (68.3%) to 4160-3930 cal BCE (68.3%) (Supplementary Fig. 22-23, Supplementary Data 2).

*Iváncsa-Lapos*

The site is located by the Danube River on a 560–570 m hill range running from North to South and protruding from the floodplain. Almost 600 features were found on an approximately 7000 $m^2$ surface area in the Southern part of the site. The vast majority of the site was comprised of the remains of an Early Bronze Age settlement. In addition to these, an 11$^{th}$-century cemetery and some features of a medieval village were excavated. An ECA single grave (Feature 335/A) was also found at the site which can be dated to 4325-4070 cal BCE (68.3%) (Supplementary Data 2).

*Polgár-Csőszhalom*

The site is situated in Northeastern Hungary, in the middle of the so-called Polgár Island in the Upper Tisza Region. The LN site belongs to the Tisza-Herpály-Csőszhalom cultural complex. The nearly 67.5 ha settlement complex is composed of at least four structural units: a tell settlement; a multiple enclosure system ringing the mound; a single-layer settlement; and another double enclosure system[9]. The first systematic excavation in the tell was started in 1957 by Ida Bognár-Kutzián and continued after 1989 by Pál Raczky. Research activities were started around the tell in 1996 by Pál Raczky and his team[10]. During the archaeological investigations at this part of site a total of 126 burials were uncovered[11]. The graves, located in various areas of the flat settlement, formed small groups consisting of two to five burials. The majority of the graves were dug in relation to houses. In the determinable cases, 30.56% of the graves were male, 43.52% of the graves were female and 25.93% of the graves were child burials[12] (Supplementary Fig. 12B). Crouching is the usual body placement, males were consistently laid on their right side, while females were positioned lying on their left side. The individuals were buried with several objects (necklaces, armrings, girdles, tools), made from various materials (*Spondylus*, bone, antler and animal teeth, stone). Bead girdles were typical pieces of the female attire, while polished stone axes were recovered from male burials[13] (Supplementary Data 2).

Based on Bayesian-modeled AMS radiocarbon measurements, the graves of the single layer settlement can be dated from 4799-4769 cal BCE to 4673-4652 cal BCE (68.3%)[14] (Supplementary Data 2).

*Polgár-Nagy-Kasziba*

The site is situated in Northeastern Hungary, in the Upper Tisza Region, in the so-called Polgár Island, near to the former Király brook. It was excavated in 1996 during the preventive excavations for the construction of the M3 motorway. Four ECA graves were found in the northern edge of the excavated area. It is not clear whether the graves belong to a larger cemetery or they form a smaller, independent grave group. Traces of a contemporaneous settlement were not detected. The graves were laid in NW-SE orientation in an extended position. They contained a large number of grave goods, Tiszapolgár-style potteries, stone and copper ornaments, stone tools and animal bones[15] (Supplementary Data 2).

Based on Bayesian-modeled AMS radiocarbon measurements, the graves can be dated from 4520-4360 cal BCE to 4335-4270 cal BCE (68.3%)[16] (Supplementary Data 2).

*Rákóczifalva-Bagi föld, Site 8*

The site is situated on the eastern bank of the Tisza River in the Middle Tisza region. A total of 4.4 ha of a multiperiod site were excavated in 2006-2007 during a preventive excavation. Traces of an ECA settlement (a building, two wells and several pits surrounded by a ditch) and nine contemporaneous burials were found. The graves were scattered throughout the surface, did not form a grave-group. A disturbed grave (F521/S610) was analyzed which was found with two vessels in relation to the pit F476/S572 (Supplementary Data 2). The grave is situated in the southern part of the excavated area. Stylistically, the pottery bears the characteristics of Tiszapolgár-Kisrétpart-style[16,17].

Based on Bayesian-modeled AMS radiocarbon measurements, the site can be dated from 4525-4380 cal BCE to 4335-4225 cal BCE (68.3%)[16] (Supplementary Data 2).

*Tiszapolgár-Basatanya*

The site is situated in Northeastern Hungary, in the Upper Tisza Region, in the so-called Polgár Island, along the Selypes creek. Ferenc Tompa conducted the first excavation in 1929 when he revealed eleven graves and two pits. Then, Ida Bognár-Kutzián excavated the site between 1950 and 1954. She revealed one LN and 154 Copper Age (CA) graves. Except for the graves destroyed by the Szandalik canal, this is considered to be the complete CA cemetery. Bognár-Kutzián distinguished period I (Tiszapolgár, ECA) and period II (Bodrogkeresztúr, Middle Copper Age) and a few graves with transitional characteristics. She built up the typochronology of the Early Copper Age and Middle Copper Age on the GHP using data deriving from this cemetery[18].

The majority of the burials follow strict rules: men were buried on their right side and women on their left side in a crouched position, usually in W-E orientation. However, among the Tiszapolgár-style graves, some individuals were buried in an extended position and a few in a contrary orientation. 43.05% of the burials were male, 35.76% were female, 19.87% were children and 1.32% were symbolic burials (Supplementary Fig. 12C). The burials contained a wide-range of grave goods. Pottery is the most frequent grave good, forming vessel sets[19]. Wealthy female burials were furnished with limestone beads as belts and copper ornaments. Wealthy male burials were furnished with Volhynian flint blades and stone mace heads. The tradition of placing boar mandibles in the graves continued from the LN and changed slightly, as now they were not explusive to male graves, and not only wild boar but also pig mandibles were used. Chipped stone tools, bone and antler tools, animal bones were also added as grave goods[18] (Supplementary Data 2).

Based on Bayesian-modeled AMS radiocarbon measurements[20], the cemetery can be dated from 4355-4270 cal BCE to 4150-3990 cal BCE (68.3%) (Supplementary Fig. 17-18, Supplementary Data 2).

*Urziceni-Vamă*

The ECA Urziceni-Vamă cemetery is located in the free zone of the Romanian-Hungarian border, on a small terrace in the marshy valley of the Negru brook. The cemetery was discovered in 2003 during construction work for duty-free shops, upon which 17 undisturbed graves and three disturbed graves were excavated. The research was extended in 2005, and the spectacular nature of the discoveries prompted much interest from specialists, leading to the resumption of the excavations beginning in 2014. In the western area, the cemetery is disturbed by a channel and the current road DN 1F. Until now almost the complete cemetery[21–23], has been investigated, presenting 132 excavated graves, containing gold jewelry and other artifacts of special value made of copper, shell, antler and stone.

Inhumation was the general burial practice. The deceased are placed in a crouched position, lying on the left or the right side, with the head facing west or east[24]. In the determinable cases, 27.88% of the burials was male, 43.27% were female, and 21.15% were children (Supplementary Fig. 12D). The majority of women are buried on their left sides and men on their right sides though there are some exceptions. Differences between the sexes continue with the funerary inventory. In the graves of women, the number of vessels is higher (five to seven vessels), and *Spondylus,* stone, copper and gold ornaments were found. In many cases, strings of *Spondylus* beads are found in the area of the pelvis, probably used for the ornamentation of clothing. As for men, the number of vessels is significantly less (one to two vessels) and are typically placed alongside stone or copper tools, namely arrowheads, blades, scrapers and borers[25,26] (Supplementary Data 2).

Based on Bayesian modeled AMS radiocarbon measurements[27], the site can be dated from 4285-4130 cal BCE to 4035-3940 cal BCE (68.3%) with a 140-300 years (68.3%) span of use (Supplementary Figs. 19-10, Supplementary Data 2).

#### Supplementary Note 2: Description of the pottery styles

The Tiszapolgár, Bodrogkeresztúr and Salcuţa archaeological cultures were primarily defined based on the typology of potteries[18,28–30]. The archaeological sites of the Tiszapolgár and Bodrogkeresztúr cultures can be found on the Great Hungarian Plain. These two cultures were considered to be chronologically consecutive and I. Bognár-Kutzián and P. Patay discussed meticulously the continuous transition from the Tiszapolgár to the Bodrogkeresztúr culture based on the typochronological examination of potteries[18,28,31]. However, the separation of these two cultures were not unproblematic and series of AMS radiocarbon dating and their Bayesian modelling showed a partial chronological overlapping between them, and their complex spatiotemporal relations[16,20]. This does not exclude the possibility that the pottery style changed so rapidly that it cannot be currently captured by radiocarbon dating. This is probably what we see in the case of Urziceni-Vamă, where the appearance of Salcuţa style elements is limited to one generation according to our current data.

The archaeological sites of the Salcuţa culture can be found in the Central Balkans and its pottery spread to Banat, Transylvania [30,32]. Similar pottery can be found on the Great Hungarian Plain as well where it is called Hunyadihalom culture[33,34] and can be dated between 3900 – 3750 cal BCE[16,20]. Primarily based on the typological analysis of the pottery, the spread of Salcuţa finds to the North from the Central Balkans was interpreted as population movement or cultural homogenization caused by the first infiltration of steppe people[28,32].

In the following, we describe the main characteristics of the potteries highlighting the differences between them (Supplementary Fig. 1.).

*Tiszapolgár pottery style*

Ida Bognár-Kutzián described meticulously the Tiszapolgár pottery based on the grave goods of the Tiszapolgár-Basatanya cemetery. Pedestalled bowls, conical cups, jars and tumblers with frequently applied plastic decorations such as pointed knobs and lugs. Impressed decorations usually consisted of impressed lentil-shaped dots organized into rows or geometric patterns[15,18,28].

*Bodrogkeresztúr pottery style*

The most characteristic forms are the so-called "milk jugs", flowerpot-shaped vessels, curved-profiled bowls, and two-handled cups. Incised geometrical motifs and net patterns filled with incrustation – sometimes combined with dots – frequently decorate the "milk jugs" and bowls[18,28,29,31,35,36]. I. Bognár-Kutzián distinguished Bodrogkeresztúr A and B based on the presence of *Scheibenhenkel* and stylistical elements of the Salcuţa culture [18,31,33].

*Salcuţa pottery style*

Usually black-burnished pottery that was frequently decorated with channelling and *Scheibenhenkel*. The most characteristic forms are shallow, compact globular bowls with short cylindrical necks, cups with two handles, pear-shaped bodies and truncated necks, the latter ornamented with wide oblique grooving. The handles go from the lip rim to the area of maximum diameter of the body, but they do not elevate over the lip [30,32,35].

### Supplementary Note 3: Notes to the genetic analyses

*Genetic diversity of the Urziceni community*

Through 'f4admix' proportions, sporadic but significant signals of Levantine PPN-related (I7130, I15618, I15619, I15623) and Caucasian HG-related genetic ancestry (I7134, I7135, I18115, I18154, I4089, I20808) had been reported at ECA Urziceni-Vamă[37].

We tested through individual $f_4$[38], qpAdm[39] and site-based qpAdm approaches for the contributions of these components to the LN-ECA diversity of the GHP, along with Eastern European hunter-gatherer (EHG) and Western European hunter-gatherer

(WHG) components and two potentially separable signals of southern origins (Jordan PPNB/Israel Natufian and ANF/Turkey Epipalaeolithic) (Supplementary Data 3-4).

Assuming normal distribution, we traced $f_4$ outliers within a given group via Z-score calculations (Supplementary Data 4). For each value in the columns EHG_WHG_f4 or Natuf_Pinarbasi_f4, the population-specific mean (μ) and standard deviation (σ) were first calculated. The Z-score calculation was performed to assess how each individual's value deviates from the population-specific mean.

In the $f_4$ scatterplot and $f_4$ tables (Supplementary Fig. 4, Supplementary Data 4) we ascertain some of the outliers from the PCA, and also the rather heterogeneous nature of the Middle Neolithic (MN) GHP and the ECA Urziceni-Vamă communities, but the genetic drift with Israel Natufian was not consistent with the signals of previous f4admix analyses[37].

*Characterization of smaller ECA sites on the GHP*

At ECA Polgár-Nagy-Kasziba and Rákóczifalva-Bagi föld site 8, both of which are characterized by Tiszapolgár-style pottery, we observed a low proportion (<11%) of HG ancestry that was dominated by WHG components (Supplementary Data 5A). This WHG dominance was similarly evident at ECA Iváncsa-Lapos and Budapest-Albertfalva-Hunyadi János út (referred to as Albertfalva), where Iváncsa-Lapos and one of the four individuals (grave 1037) at Albertfalva showed over 11% combined EHG+WHG ancestry, significantly deviating from the rest of the site (Z-score = 3.42, Supplementary Data 5C).

*Family structures in the Late Neolithic Great Hungarian Plain*

From the 224 graves of the LN Aszód-Papi földek site, we produced data from two males and 20 females, which is not representative of the overall sex ratio at the site (Supplementary Fig. 12A, Supplementary Data 2, see Methods). We detected only two kinship relations within the Aszód-Papi földek site (graves 127 - 128, and 56 - 57 respectively). Burials 127 and 128 were siblings. Female individuals in graves 56 and 57 were in a second degree relation. We propose two pedigree variants on Supplementary Fig. 15 based on the fact that individuals shared mitochondrial haplogroup X2d1 and had avuncular 1 or 2 relation in the form of aunt-niece relationship. Graves 56 and 57 were found next to each other in the grave group D, and Graves 127 and 128 in the grave group B[1]. Although further graves located close to each other were analyzed from grave groups A-F, no further biological relations were detected. Only seven individuals were suitable for IBD analysis. These individuals primarily exhibit between-module connections, with loose ties to both LN Polgár-Csőszhalom and ECA Tiszapolgár-Basatanya communities, via different nodes, and do not form a separate Leiden cluster ($k$=0.014, Supplementary Fig. 6).

Three kindreds/pedigrees could be identified at Polgár-Csőszhalom through first-degree relationships, including two pairs of adult-aged sisters and a couple with a young adult son (Supplementary Fig. 9). Among these, the distant connection of the three pedigrees is highlighted via grave 421 (Fig. 4B).

*Detailed connections between Polgár-Csőszhalom and Basatanya communities*

Three males (graves 406, 618, 921) and one female (grave 852) at Polgár-Csőszhalom are connected to Tiszapolgár-Basatanya via at least 2x12 cM IBD connections, suggesting local continuity in the LN-ECA transitional period. Interestingly, these are not the latest graves among the sampled ones. At ECA Tiszapolgár-Basatanya, female individuals from graves 33, 57, and 59 establish IBD connection with earlier Polgár-Csőszhalom (Fig. 5), though their maternal lines do not match their IBD pairs. Notably, the male in grave 406 at Polgár-Csőszhalom is linked to Tiszapolgár-Basatanya through grave 59 (as shown in Fig. 4B). This male carried the H46 mitochondrial DNA (mtDNA) lineage, also found in the female from grave 76 at Tiszapolgár-Basatanya, who is third or fourth degree related to grave 59 at the same site. H46 haplotype, along with other maternal connections via K2, N1a, T2b, U4 haplogroups and the differences between maternal and paternal haplogroup matches suggest that the connection between the two sites may have been predominantly maternal (Supplementary Fig. 10A-B). Paternal lineages are more diverging, Polgár-Csőszhalom exhibiting G2a and J2a, whereas Tiszapolgár-Basatanya having C1a, beside various I2a lineages among shared Y-chromosomal types (Supplementary Fig. 11). Nevertheless, direct matches between the uniparental connections/similarities and the IBD connections were not found in the dataset.

*Kinship structures and individual observations at Tiszapolgár-Basatanya*

At Basatanya, we observed a significant difference in runs of homozygosity (ROH) between the two siblings in graves 149 and 156, with grave 149 showing evidence of close parental consanguinity (likely third-degree related, such as first cousins, as indicated in Supplementary Fig. 5). Simulations have shown considerable variability in ROH patterns among offspring of first cousins[40], suggesting that while grave 156 lacks ROH blocks >20 cM, this sibling was still the product of close relatives. Neither the IBD nor the ROH results for the individual in grave 149 revealed any chromosomal abnormalities, which might otherwise explain the notable difference in ROH patterns between the siblings.

Grave 23 of the Family A in the Tiszapolgár-Basatanya cemetery is one of the richest male burials, reflecting the continuity of LN burial traditions, including prestige and status goods indicative of his wealth and rank (stone macehead, copper ornaments, wild boar mandible, Volhynian flint blades). This individual was second-degree related on the paternal side to the male in Grave 28, and more distantly related to his descendants. In Grave 52, a man who was probably the son of first-cousins or 3rd-degree relatives (Supplementary Fig. 5A), with similarly rich grave goods, is 3rd/4th-degree related to Grave 23. However, another branch of Family A shows a more nuanced picture: the male in Grave 61, who also exhibits signs of being an offspring of first-cousin marriage, was not as richly buried as the individual in Grave 52.

Comparing the ROH signals with pottery styles of the grave inventories at Basatanya, we observe that high levels of homozygosity (between 20 and 161 cM sum total of ROH segments >20 cM) were not restricted to individuals buried with a certain style, as we detected ROH from three Bodrogkeresztúr- and five Tiszapolgár- and in one transitional-style graves ($p=0.66$ in permutation test, Supplementary Data 1 and 6).

Outliers in their genetic ancestry compositions are limited at Basatanya. Two were identified as PCA outliers (graves 21, 145), two as IBD outliers (graves 45 and 83) and only three individuals failed at the ANF-WHG-EHG 3-way qpAdm test (Fig. 2, Supplementary Data 5 and 7). One female individual (I5101/21) from a Tiszapolgár-style grave displayed over 15% WHG ancestry (ca. 16.8±2.9% on qpAdm, Z-score=3.82 for the deviation from the mean WHG component in her group) (Fig. 2A, Supplementary Data 5C). In sharp contrast, another female individual with Bodrogkeresztúr-style pottery (I8796/145) exhibits as low as 6.3% total HG ancestry—a notable deviation from her group, with a combined HG ancestry Z-score of -3.45 (Supplementary Data 5C).

*Connection system of the Neolithic and Copper Age Urziceni communities*

The MN Piscolț phase of Urziceni-Vamă is represented by two individuals, who showed 81-86% ANF with 13-16% WHG and only negligible EHG ancestry in qpAdm analyses (Supplementary Data 5). The high level of WHG ancestry observed (Z-score=1.46-2.11, considering the average WHG in the whole LN-ECA dataset as μ) during the MN period decreased in later periods at the site, where 28 out of 50 ECA individuals could still be modeled using a local MN source alone (Supplementary Data 5Eb).

A Neolithic individual from Urziceni-Vamă (grave 5) shows LN GHP IBD connections, similar to the male from grave 81, whose failed radiocarbon dating could not ascertain his dating.

Two females in a mother-child relationship (graves 18 and 46) entered the community from external groups. Individual from grave 18 from Urziceni is only connected to Tiszapolgár-Basatanya grave 66, while grave 46 has more within-cemetery IBD connections, highlighting an asymmetrical connection system (Fig. 4).

A total of 44 distinct mtDNA subhaplogroups were described (Supplementary Fig. 10) at Urziceni, including haplogroups pointing toward the southeast, such as N1b, and others (e.g. J2b1, J1c) with parallels to Varna and Pietrele sites[41]. Further haplogroups point toward the east or hint at traces of HG ancestry[42] (basal U4, U5a) among typical groups of local Neolithic origin (different H-s, K1a, K2a, N1a, U4, T2(b), T2e, and W1[43–45]. The embeddedness of these later haplotypes are also visible on the median joining network of the main CB sites' maternal lineages (Supplementary Fig. 10B).

If we compare these lineages with the PCA outliers, J1c emerges in four cases as a potential external lineage (three males and one female) in the community. However, its prevalence in the LN Aszód-Papi földek community (Supplementary Fig. 10) and also the diverse genetic composition of these outliers suggest that conclusions drawn from maternal lineages should be approached with caution.

The paternal genetic diversity at Urziceni-Vamă include C2a of Mesolithic and H2 and G2a elements of Neolithic origin[43], as well as potentially new external influences such as R1b from steppe-outliers (graves 12 and 79, Supplementary Fig. 11). The presence of J2a in graves 11 and 59 may indicate southeastern connections[41], though this lineage was also detected in MN Rákóczifalva–Bagi-földek and LN Polgár-Csőszhalom[46].

Consanguinity was rare in the Urziceni-Vamă cemetery, with only two cases observed via ROH analysis. One was a woman buried with gold jewelry (Grave 29), for whom no relatives were found in the cemetery, suggesting she may have joined the community as an adult. The other case involved a male in Grave 71 with common grave goods, distantly related to a woman in Grave 97. At Urziceni-Vamă, pottery styles do not show a correlation with signs of inbreeding either, leading us to conclude that the practice of consanguineous marriages was not associated with distinct pottery stylistic traditions of the sampled communities. It is however noteworthy that the outlier individuals observed on PCA bear no or only limited (4-8 cM) ROH signals.

### Supplementary Figures

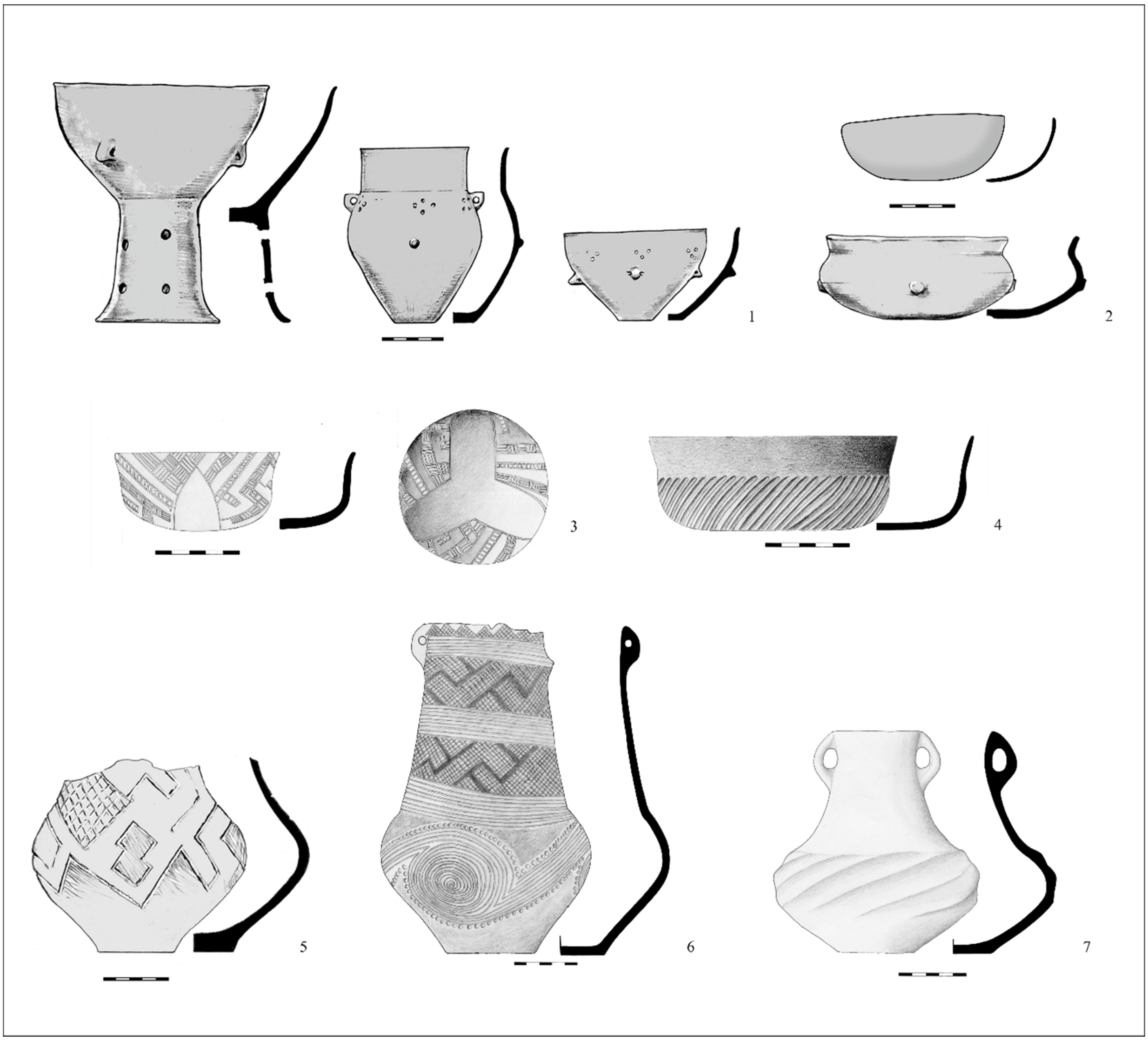

**Supplementary Figure 1.** Tiszapolgár-style (1) and Bodrogkeresztúr-style (2, 5) potteries from the Tiszapolgár-Basatanya cemetery (modified after Raczky & Siklósi 2013[20], figures 3 and 4), Bodrogkeresztúr-style (3, 6) potteries and Bodrogkeresztúr-Salcuţa-style potteries (4, 7) from the Urziceni-Vamă cemetery.

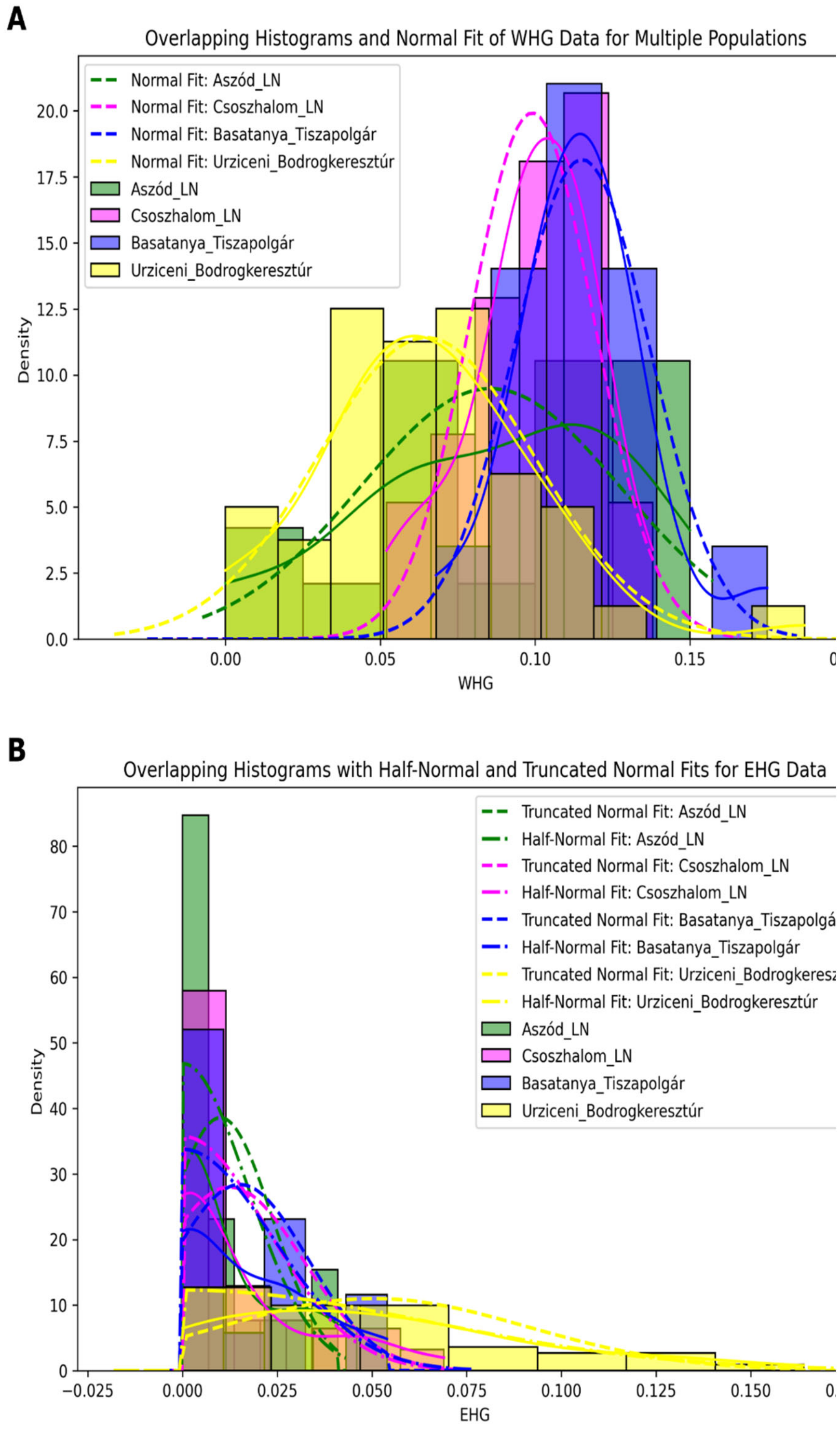

**Supplementary Figure 2. Histograms of WHG (A) and EHG (B) qpAdm ancestry components, calculated in 3-way models for each sample with fitted normal probability density function.**

Plot A shows individual WHG components (shown as density plots) are normally distributed in the groups. The EHG component presented on plot B however, is half-normal or truncated normally distributed, as many individuals have zero EHG ancestry. Individual qpAdm data are presented in Supplementary Data 5B. Histograms were generated using the sns.histplot function of Seaborn library in Python 3.12. Note that the height of the bars shows the density per bin width, and not the raw count of the data points. Script for plotting can be found at GitHub (github.com/ArchGenIn/Szecsenyi-Nagy_2025) and at Zenodo (DOI: 10.5281 /zenodo.15221967).

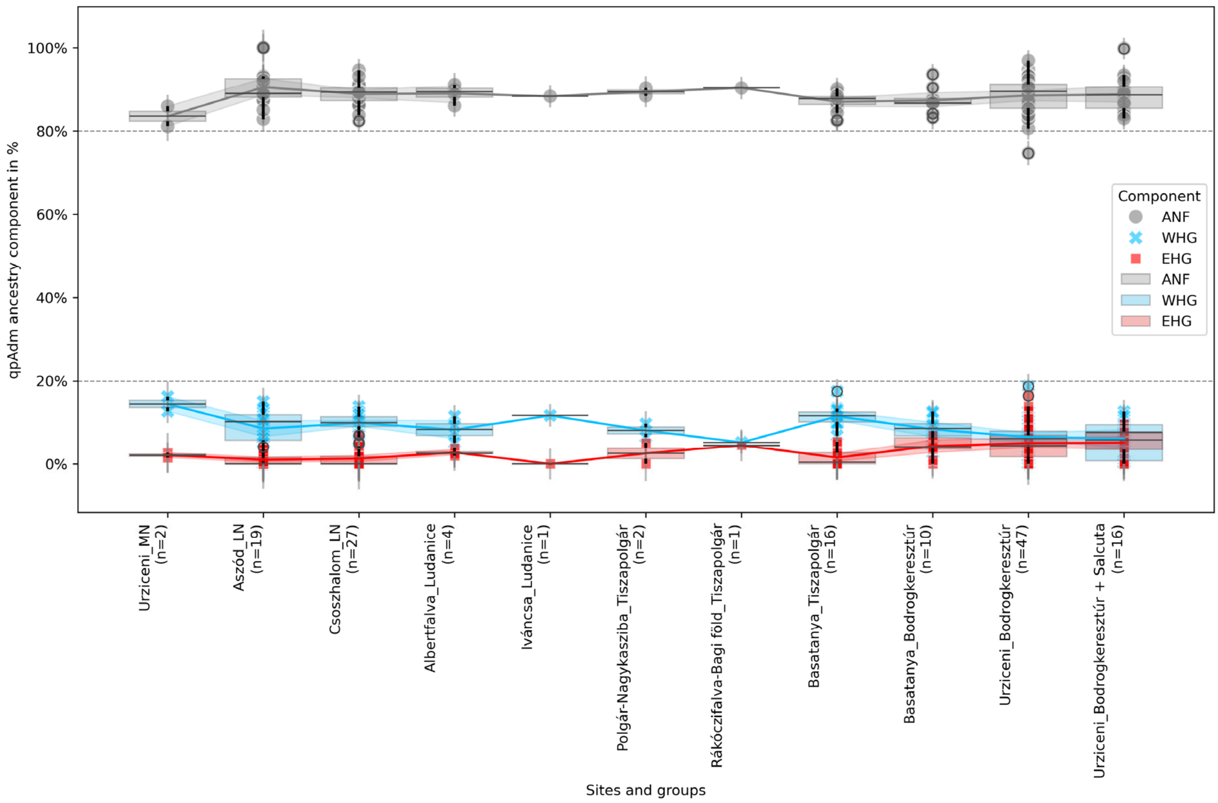

**Supplementary Figure 3. Individual qpAdm 3-way models presented as boxplot per site and pottery styles.**

Abbreviations in the legend: WHG: Western Hunter-Gatherer, EHG: Eastern Hunter-Gatherer, ANF: Anatolian N ancestry component. Source data of the box plot elements (centre line that represents the median of the data, box limits that correspond to upper (Q3) and lower (Q1) quartiles; the interquartile range (IQR), whiskers (black lines) are provided in the Source Data file. Points treated beyond the whiskers are treated as outliers (marked with black circles), also listed in the Source Data file. The sample size of a given group and individual qpAdm results are shown in Supplementary Data 5a. Script for plotting can be found at GitHub (github.com/ArchGenIn/Szecsenyi-Nagy_2025) and at Zenodo (DOI: 10.5281 /zenodo.15221967).

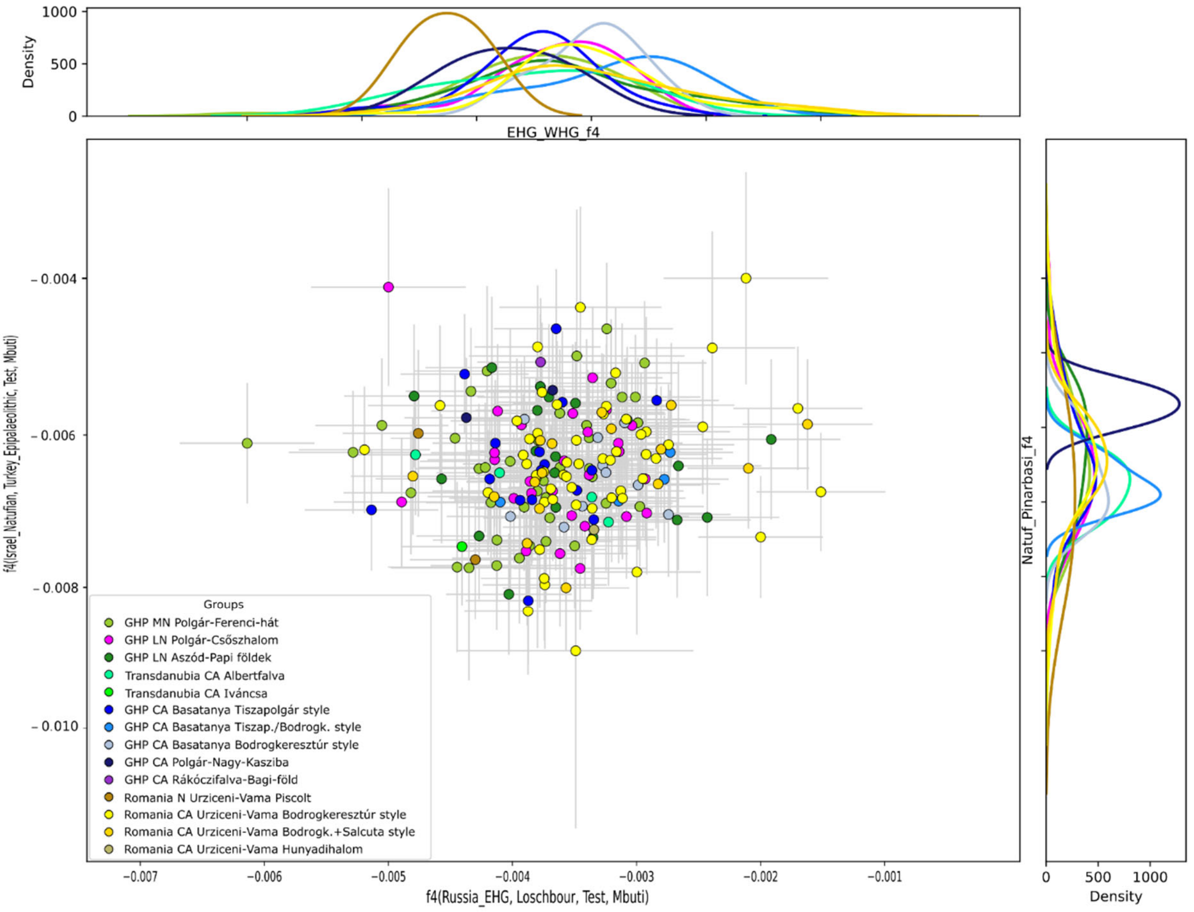

**Supplementary Figure 4. Scatterplot of $f_4$ values with standard errors, calculated for genomes of this study and comparative data, along with the density plots of the two $f_4$-statistics.**

We calculated $f_4$-statistics in two forms: $f_4$(Israel_Nautfian, Turkey_Epipalaeolithic, Test, Mbuti) plotted along y axis and $f_4$(Russia_EHG, Loschbour, Test, Mbuti) plotted along x axis. Individual $f_4$ data with Z-scores and are seen in Table S3. All samples share more genetic drift with the Anatolian Epipalaeolithic than with the Israel Natufian and with Loschbour (WHG) than with Russia EHG. We identified some of the outliers from the PCA (Fig. 2A), as well as the rather heterogeneous nature of the MN GHP and the ECA Urziceni-Vamă communities. We detect variation of allele sharing with Levantine populations, but allele sharing with Anatolian Epipalaeolithic is significantly more in every case and the fluctuation cannot be matched to any chronological or geographical trends, as it was already present in the Early Neolithic of the study area (Supplementary Data 4, Lazaridis et al.[37]).

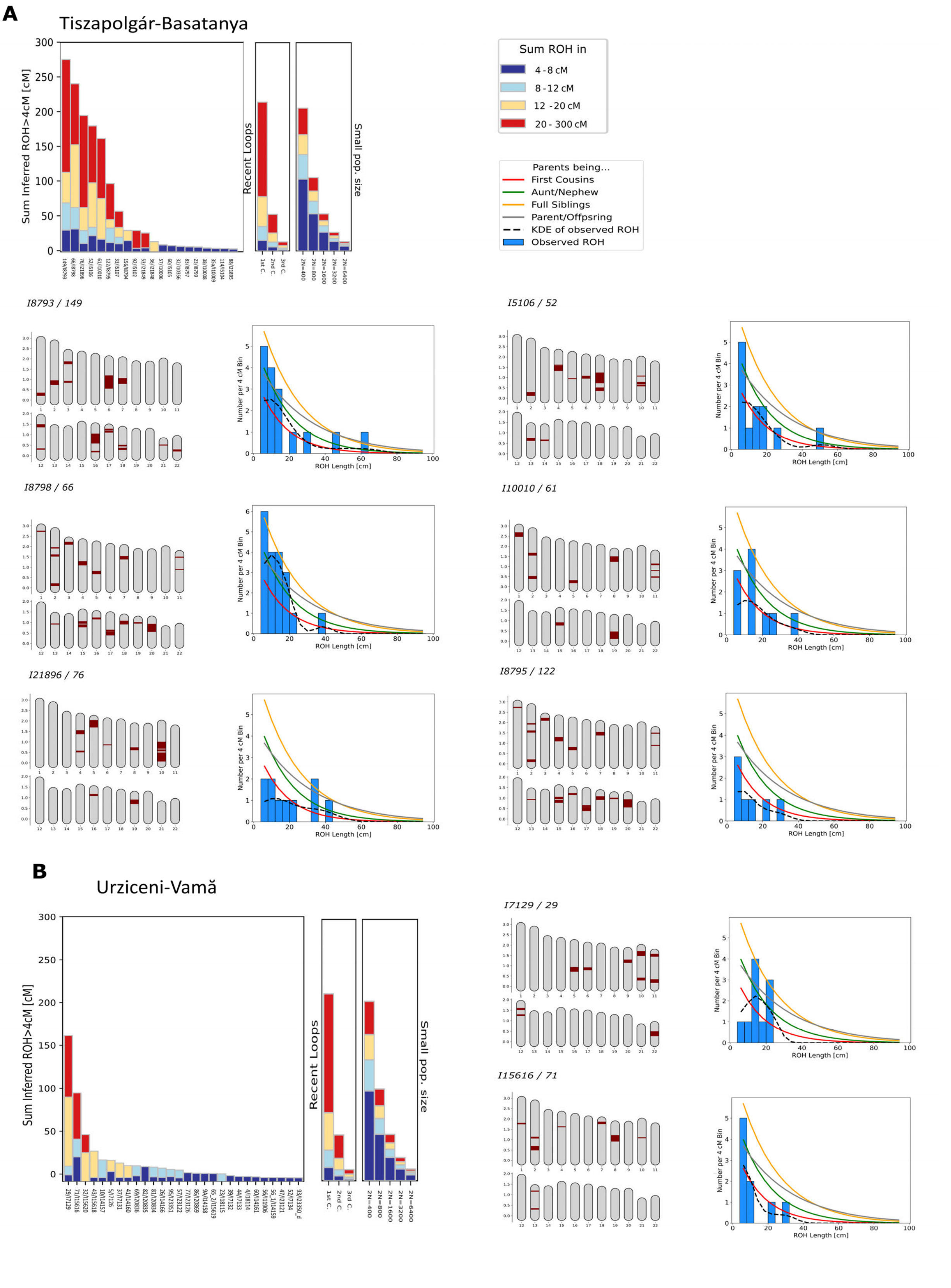

**Supplementary Figure 5. Individual ROH pattern of the inbred individuals from site Tiszapolgár-Basatanya (A) and Urziceni-Vamă (B)**

On the barplots, each individual is represented by stacked vertical bars, where the length of each bar is determined by the ROH of this individual falling into four length classes (4–8, 8–12, 12–20, and >20 cM, color-coded). The position of the ROH is marked on the 22 autosomes (maroon), with map length annotated in Morgan. We depict a histogram of the ROH lengths, together with expected densities of ROH for certain degrees of parental relationships, calculated and presented as described in Ringbauer et al.[40]. Kernel density estimation (KDE) of the ROH distribution is also shown on each histogram. The individual histograms depict individuals who were probably offsprings of 3rd degree relatives. In graves Basatanya 33, Urziceni-Vamă 32, Polgár-Nagy-Kasziba 38 and Albertfalva 1037 probably children of 4-5th degree relatives (e.g. 2nd cousins) were buried. Source data of the bar charts are provided as a Source Data file.

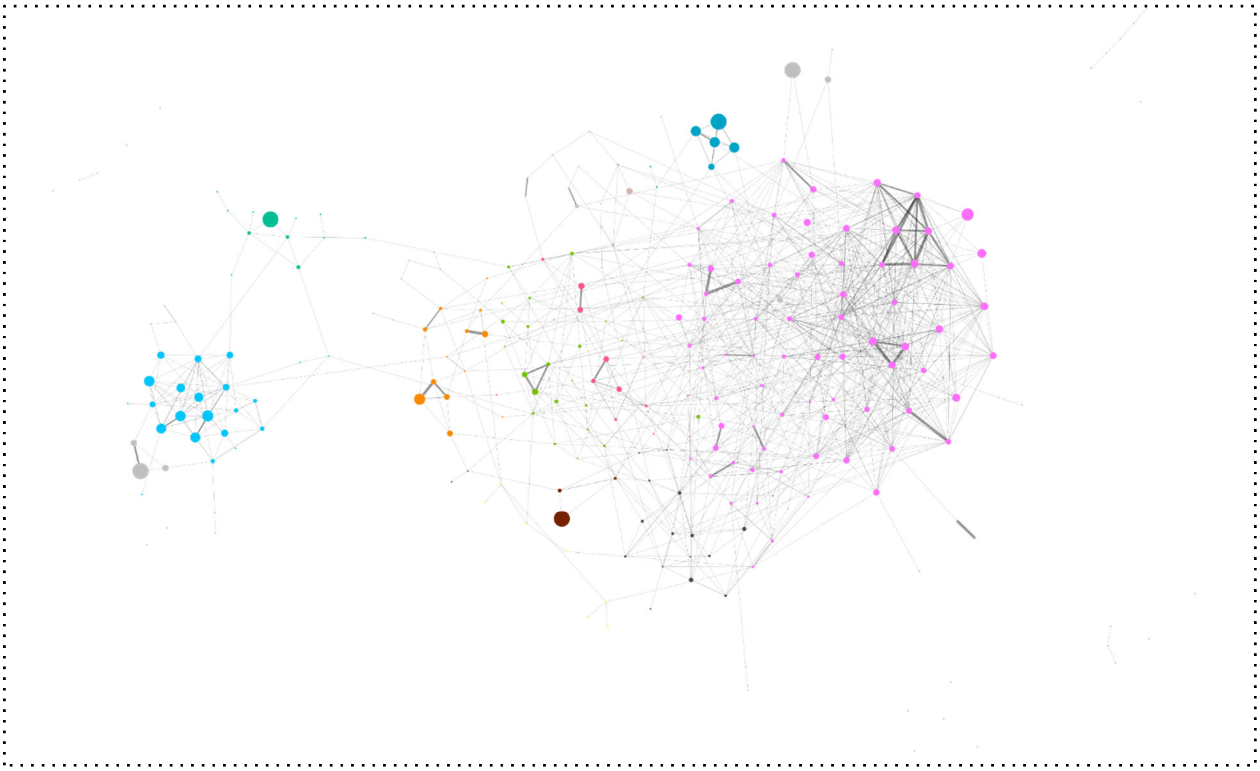

**Supplementary Figure 6. Leiden community detection and cluster coefficient with min. 1x12cM edges considered.**

Node size in the network graph is proportional to the clustering coefficient, which ranges from 0 to 0.73, with smaller circles representing lower coefficients. This metric indicates how interconnected an individual's neighbors are. The visualization corresponds to Figure 4A in the main text. For community detection, the Leiden algorithm was used with a resolution of 0.04, based on the constant Potts model[47], which partitioned the graph into 52 clusters with a quality score of 0.77. The top 10 largest clusters are highlighted with distinct colors, and edges are weighted according to total IBD (identity by descent) sharing. It is seen on the plot that the largest, pink cluster includes 27 individuals from Basatanya, 25 from Polgár-Csőszhalom, one from Aszód and two from Polgár-Nagy-Kasziba, as well as 19 individuals from Urziceni. Most individuals from Urziceni display low clustering coefficients (0.178 on average, compared to 0.33 at Basatanya and 0.216 at Basatanya in the same cluster), except for those who are part of nuclear families (Supplementary Data 7). The Leiden algorithm is an improved method for detecting modular structures in networks, building on the Louvain algorithm. Its main limitation is its assumption of non-overlapping communities, which can fail to capture complex, interlinked group structures in datasets with overlapping populations. However, since we did not use Leiden to argue for separating potentially overlapping communities, this limitation does not undermine our results. Source data of the network are provided as a Source Data file and Supplementary Data 7.

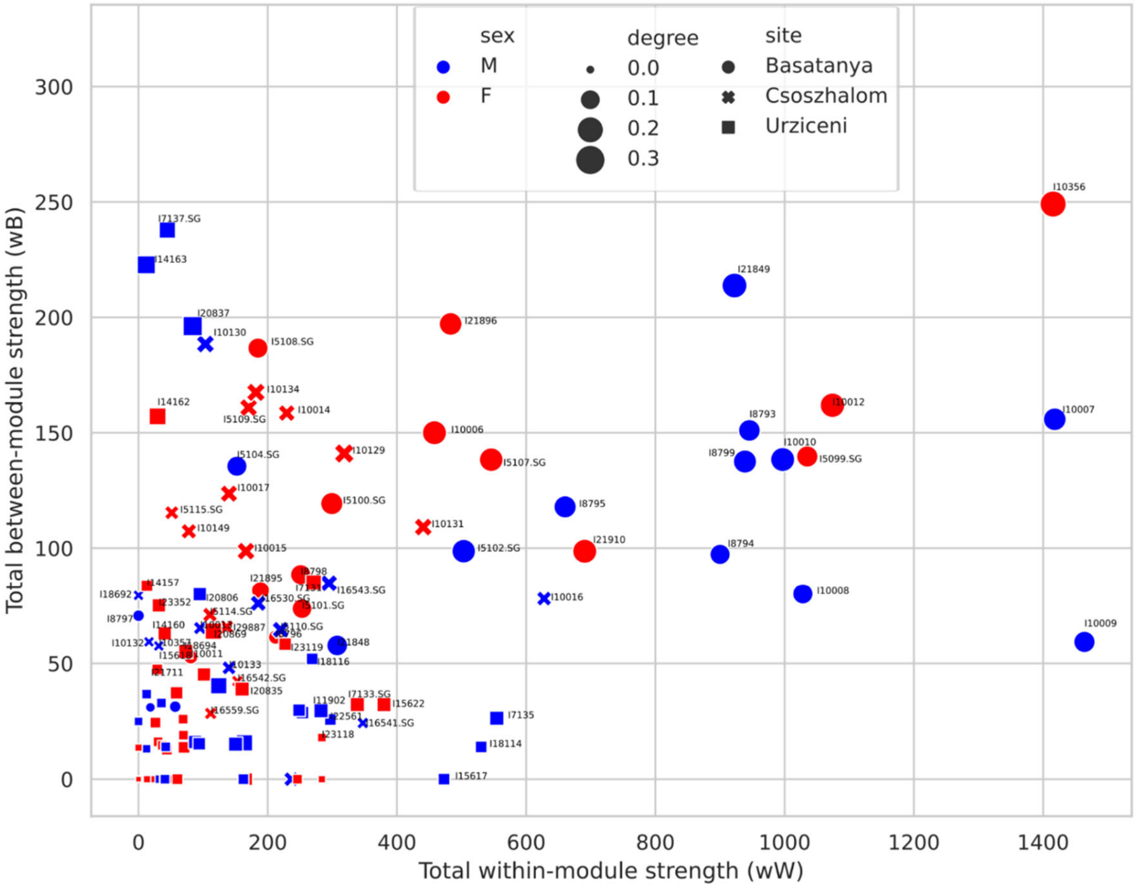

**Supplementary Figure 7. Scatterplot of accumulated within-module vs. accumulated between-module strength (w), for the subgraph of the three sites Tiszapolgár-Basatanya, Polgár-Csőszhalom and Urziceni-Vamă.**

At least 12 cM IBD connections were the basis of the calculation. Maximum IBD lengths were used as weights of the links attached to each node, and these weights were summed separately for links within the same site ("within strength") and for links to other sites ("between strength"). Weights are not normalized by the maximum length/sum IBD, but differentiated between and within the modules (the sites). Degree means degree centrality (*k*), defined as the number of links held by the node in the graph. Exact numbers are seen in Supplementary Data 8C. This plot demonstrates that members of the Basatanya community exhibit the strongest within-module connections and the most connections per individual, while some also maintain strong connections to other sites. Script for plotting can be found at GitHub (github.com/ArchGenIn/Szecsenyi-Nagy_2025) and at Zenodo (DOI: 10.5281 /zenodo.15221967).

**A**

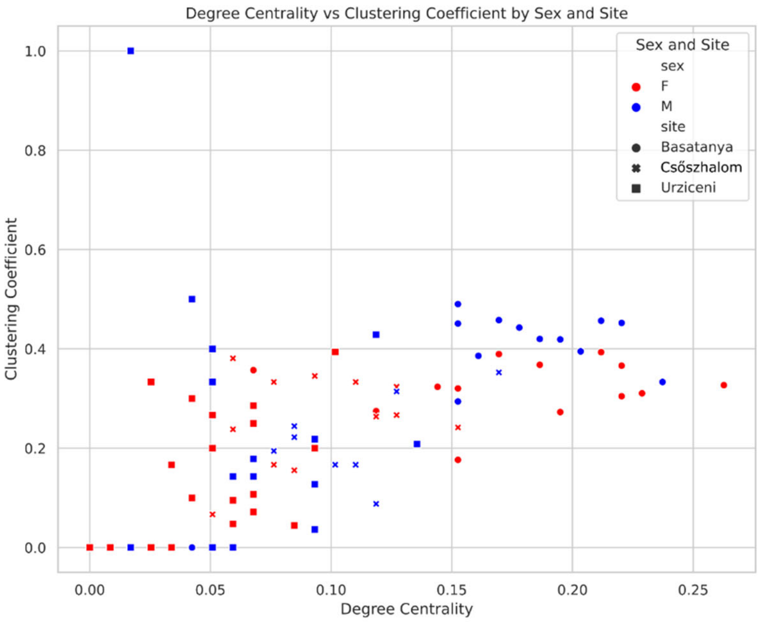

**B**

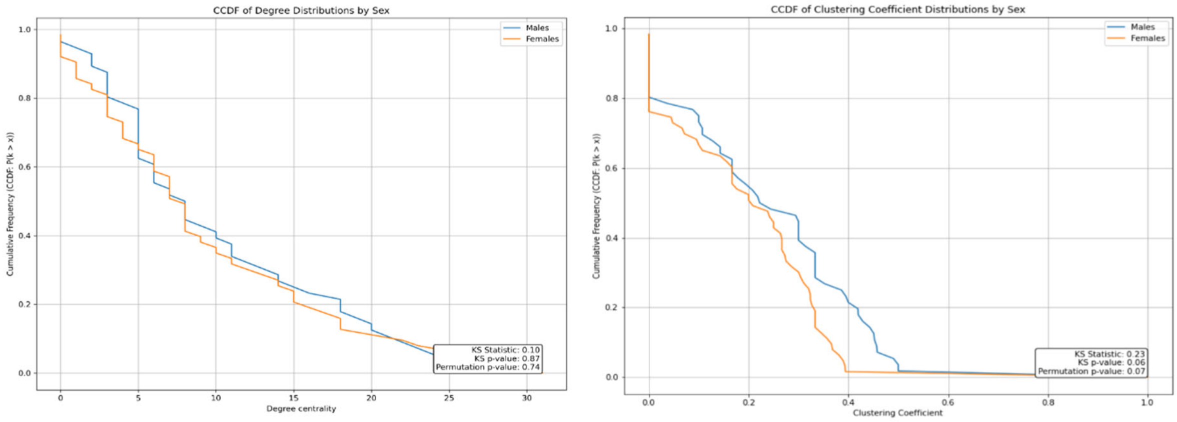

**Supplementary Figure 8. A: Scatterplot showing degree centrality versus clustering coefficient by sex and sites, and distributions of these parameters per sex. B: Distributions of degree centrality and clustering coefficient of males and females at all three sites considered together.**

At least 12 cM IBD connections were used as the basis for the calculations in the NetworkX package in Python. Degree centrality (k) for each node was computed as the number of links to all other nodes, then normalized using NetworkX's degree_centrality function, which divides the raw degree by (N − 1) where N is the total number of nodes in the graph (Supplementary Data 8). The clustering coefficient for each node in a graph (represented on y-axis) is a measure of how interconnected a node's neighbors are. Higher clustering coefficient was observed among males compared to females (Supplementar Data 7). Nevertheless, the distributions of degree centrality and clustering coefficient of males and females were not significantly different according to two-sample Kolmogorov–Smirnov tests (panel B). Here, CCDF (complementary cumulative density function) refers to the probability that $k \geq X$. The permutation-based KS tests ($n = 1000$) yielded $p > 0.05$, indicating no statistical difference. Scripts for statistical calculations and plotting can be found at GitHub (github.com/ArchGenIn/Szecsenyi-Nagy_2025) and at Zenodo (DOI: 10.5281 /zenodo.15221967).

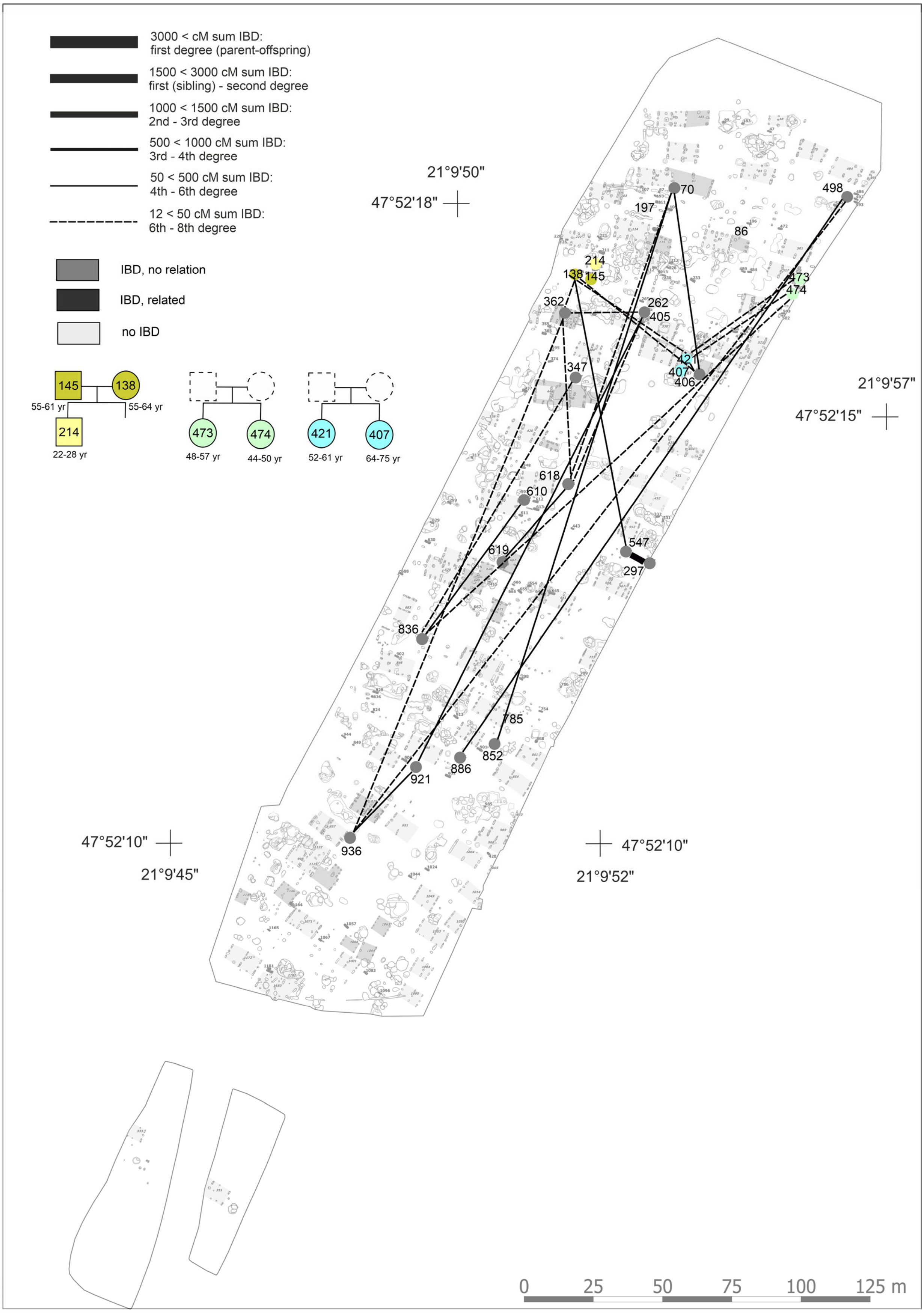

**Supplementary Figure 9. IBD connections projected on the map of the Polgár-Csőszhalom Late Neolithic site with genetic kindreds.** Source data are provided as a Source Data file, kinship is presented in Supplementary Data 12.

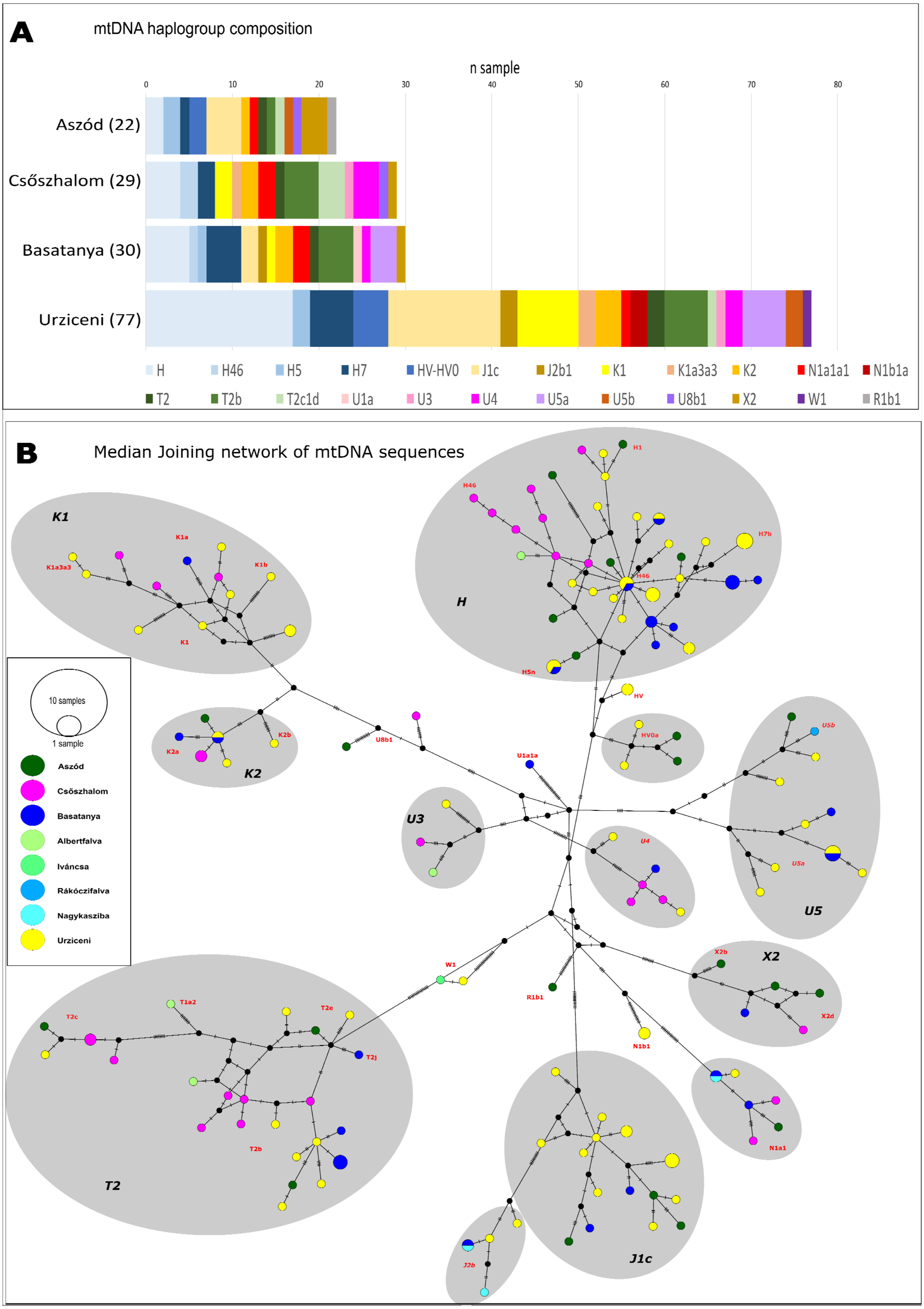
A
mtDNA haplogroup composition
n sample
0
10
20
30
40
50
60
70
80
Aszód (22)
Csőszhalom (29)
Basatanya (30)
Urziceni (77)
H
H46
H5
H7
HV-HV0
J1c
J2b1
K1
K1a3a3
K2
N1a1a1
N1b1a
T2
T2b
T2c1d
U1a
U3
U4
U5a
U5b
U8b1
X2
W1
R1b1
B
Median Joining network of mtDNA sequences
K1
K1a
K1b
K1a3a3
K1
K2b
K2a
K2
U8b1
H
H1
H46
H7b
H5n
HV
HV0a
U1a1a
U3
U4
U5b
U5a
U5
X2b
X2
X2d
W1
R1b1
N1b1
N1a1
T1a2
T2c
T2e
T2j
T2b
T2
J1c
J2b
10 samples
1 sample
Aszód
Csőszhalom
Basatanya
Albertfalva
Iváncsa
Rákóczifalva
Nagykasziba
Urziceni

**Supplementary Figure 10. Comparison of the maternal genetic diversity at Late Neolithic Aszód-Papi földek, Polgár-Csőszhalom and Early Copper Age Tiszapolgár-Basatanya and Urziceni-Vamă sites.**

Subfigure A: MtDNA haplogroup distribution. Haplogroups given in absolute numbers, were determined by HaploGrep (see Methods). Some subgroups are specified as they show connections between some of the sites under study. Details are presented in Supplementary Data 1. Subfigure B: Median joining network of the mtDNA sequences from the listed LN-ECA sites.

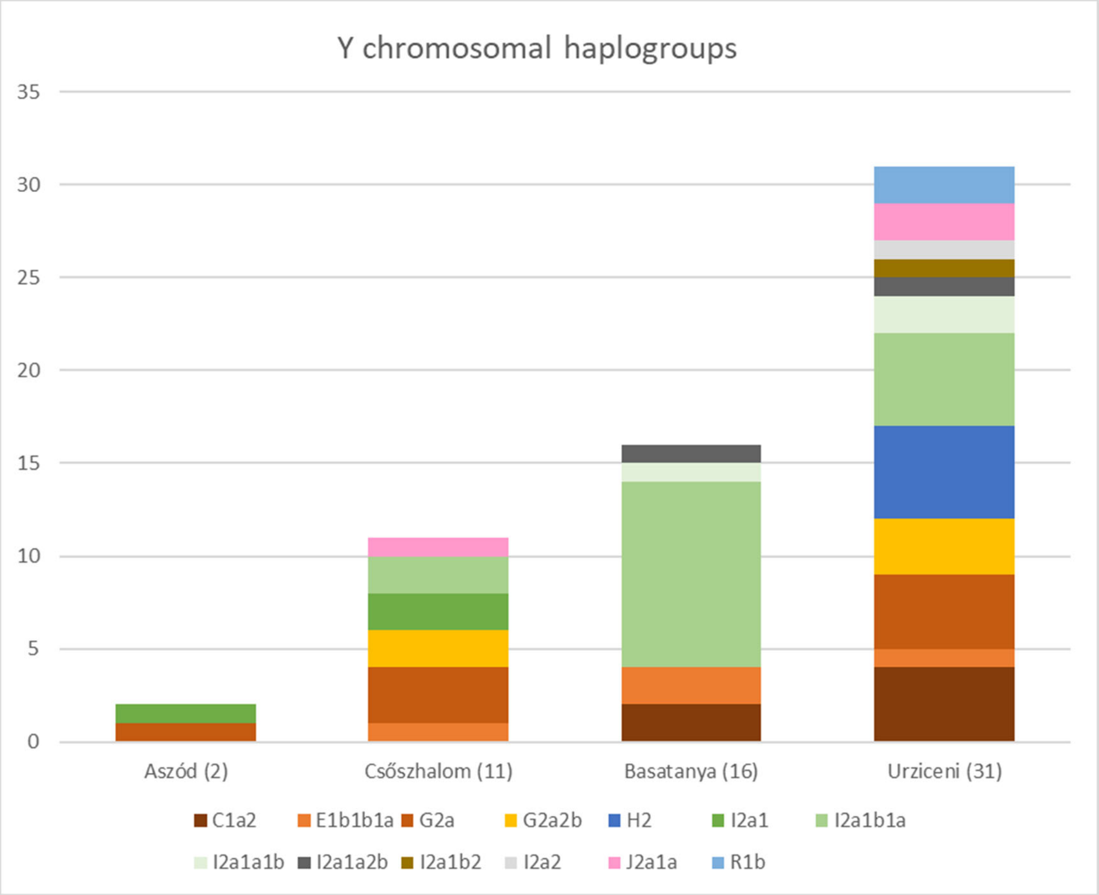

**Supplementary Figure 11. Comparison of the paternal genetic diversity at Late Neolithic Aszód-Papi földek and Polgár-Csőszhalom and Early Copper Age Tiszapolgár-Basatanya and Urziceni-Vamă sites.**

Data are given in absolute numbers for males. Y haplogroups were determined with the Yleaf program (see Methods). Resolution of within haplogroup diversity depends on sample preservation, therefore we note that some samples in categories such as G2a or I2a1 might belong to subgroups of these. Details are presented in Supplementary Data 1.

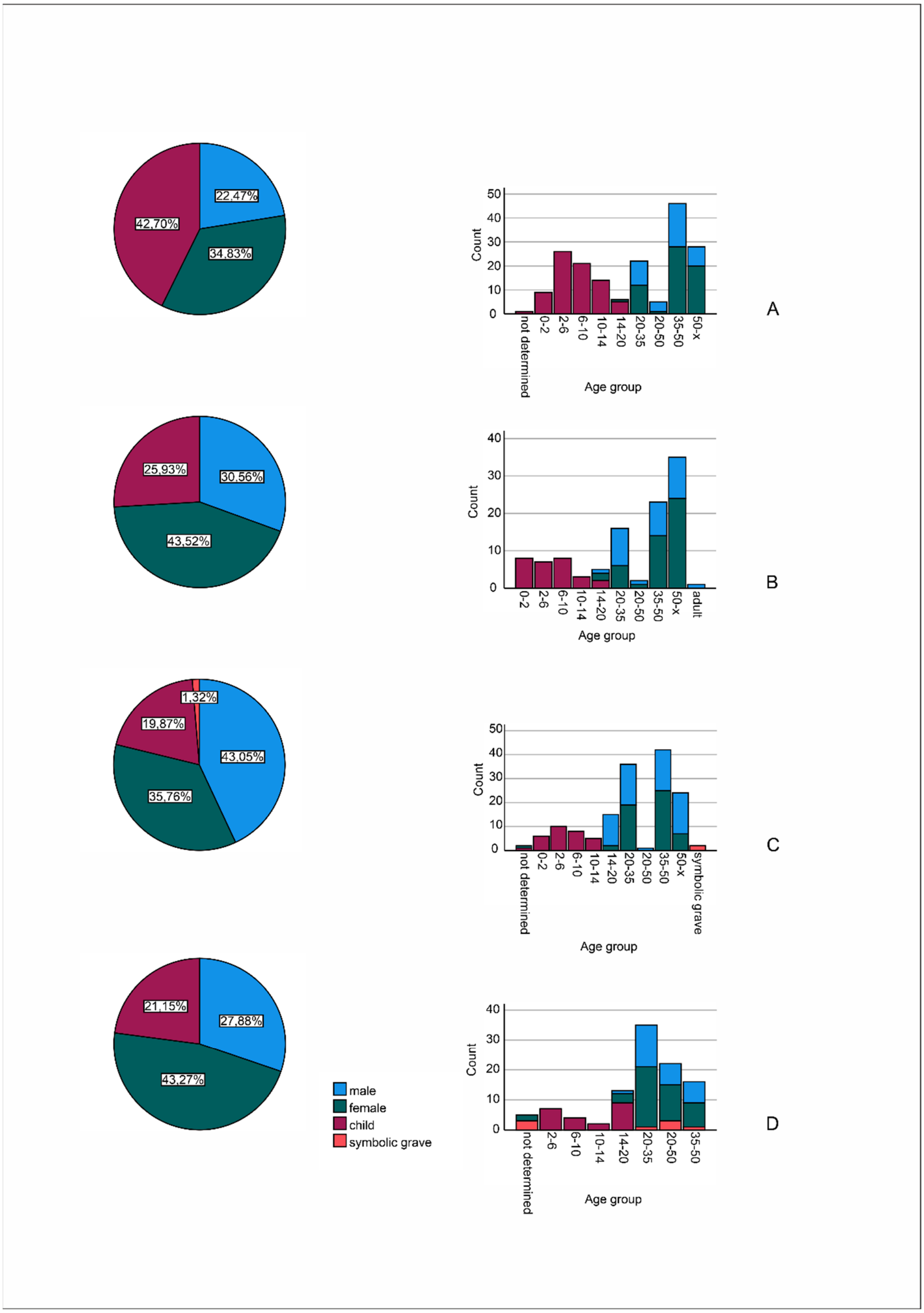

**Supplementary Figure 12. Frequency of male, female, child graves and age categories in Aszód-Papi földek (A), Polgár-Csőszhalom (B), Tiszapolgár-Basatanya (C) and Urziceni-Vamă (D) sites based on the anthropological examination.** Background data are provided as a Source Data file.

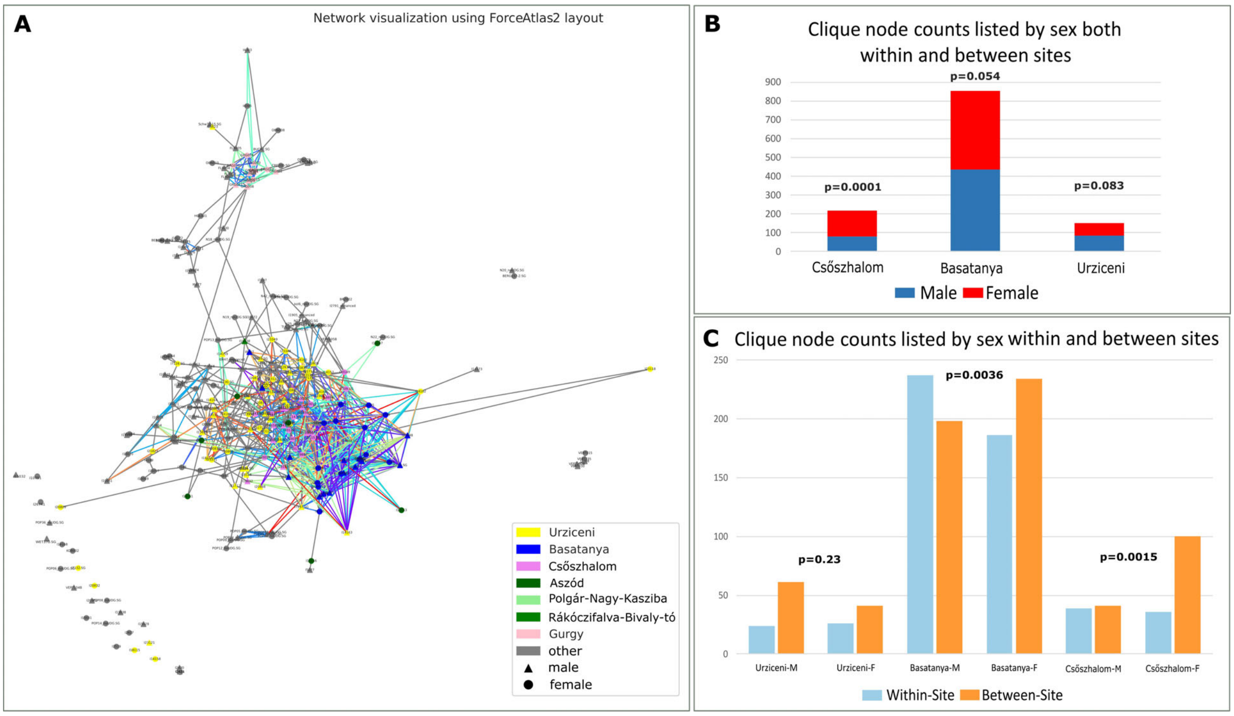

**Supplementary Figure 13. Network cliques and their statistics.**

Clique is measured within the network as subsets of at least three nodes, where each pair of nodes is adjacent, forming a complete subgraph. Subfigure A: IBD graph using ForceAtlas2 layout highlighting the cliques with colored edges. If an edge belongs to multiple cliques, it is colored based on the first clique it is found in. The network was drawn in NetworkX and fa2 packages in Python 3.7. Cliques themselves are presented in Supplementary Data 9. Subfigure B: Sex distribution of the cliques and Fisher's exact test p values. If a clique spans multiple sites, it appears in the counts for each of those sites. It is seen on this graph that the participation of the females were significantly more in cliques than that of the males at Csőszhalom. Subfigure C: Distribution of cliques within and between the sites, considering both sexes. The two sexes shared significantly different proportions within and between site cliques at Csőszhalom and Basatanya. The Chi-Square test was calculated for each site-sex group and their p-values are reported in the chart (exact data and statistics are shown in Supplementary Data 8B, code is available at at GitHub (github.com/ArchGenIn/Szecsenyi-Nagy_2025)).

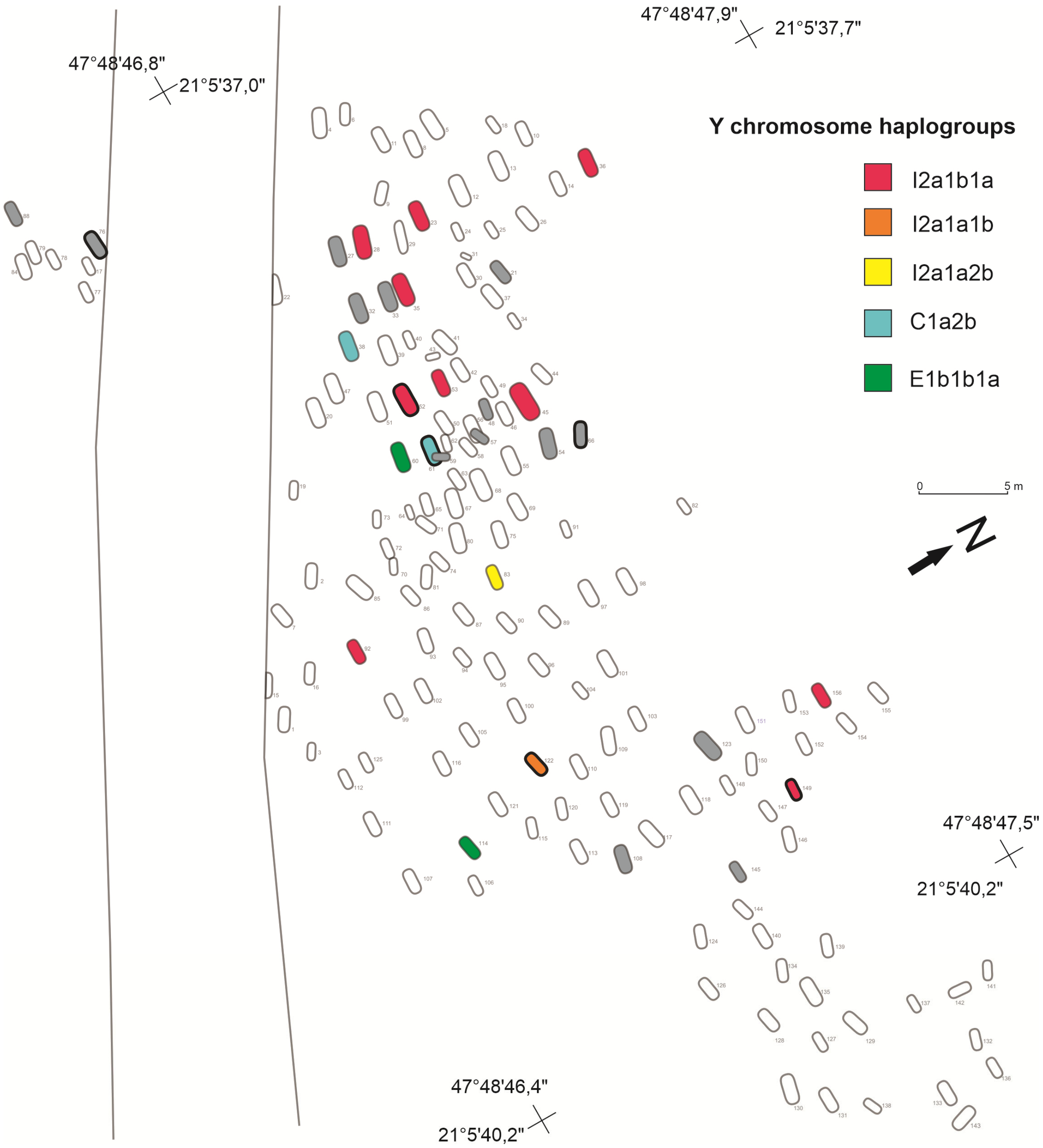

**Supplementary Figure 14. Distribution of paternal lineages in the ECA Tiszapolgár-Basatanya cemetery.**

Haplogroup data per sample and grave is shown in Supplementary Data 1.

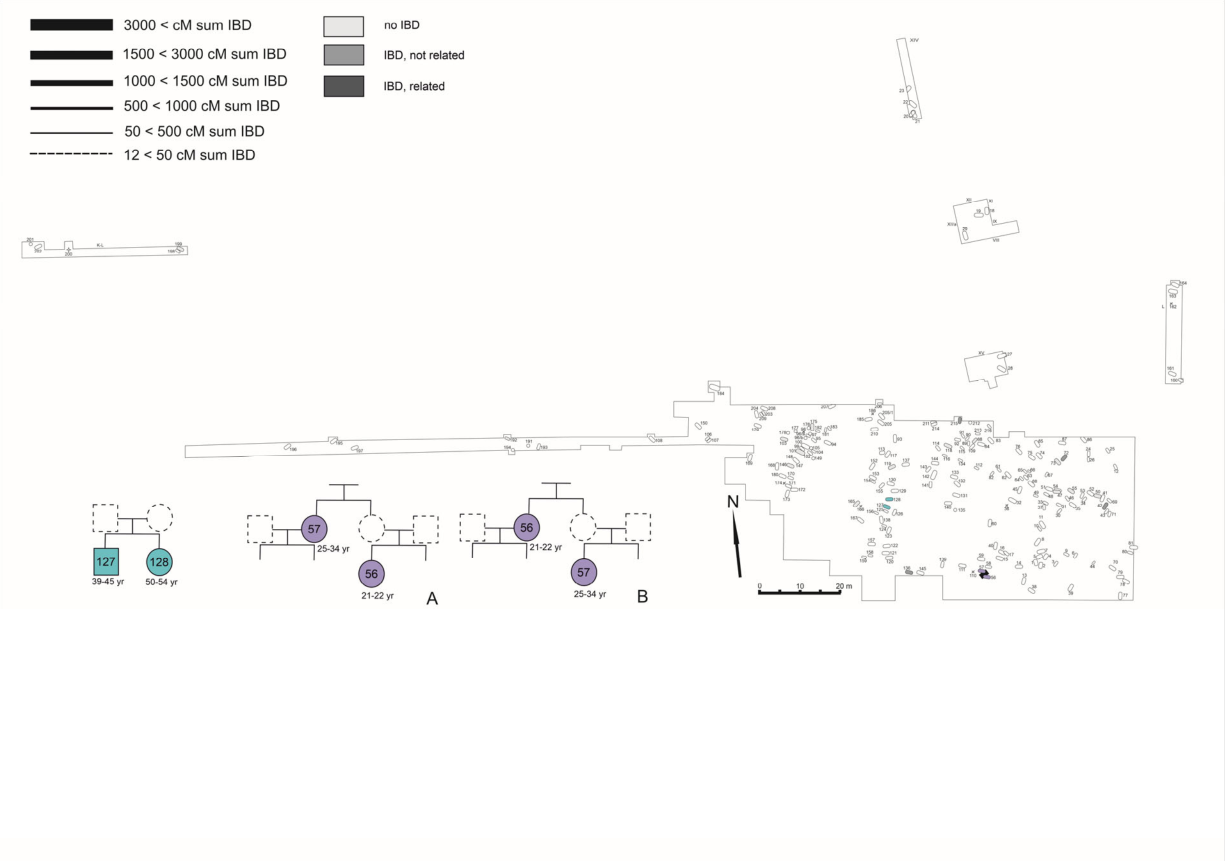

**Supplementary Figure 15. Map of Late Neolithic Aszód-Papi földek burials and their biological relatedness.**
Source data are provided as a Source Data file, kinship is presented in Supplementary Data 12.

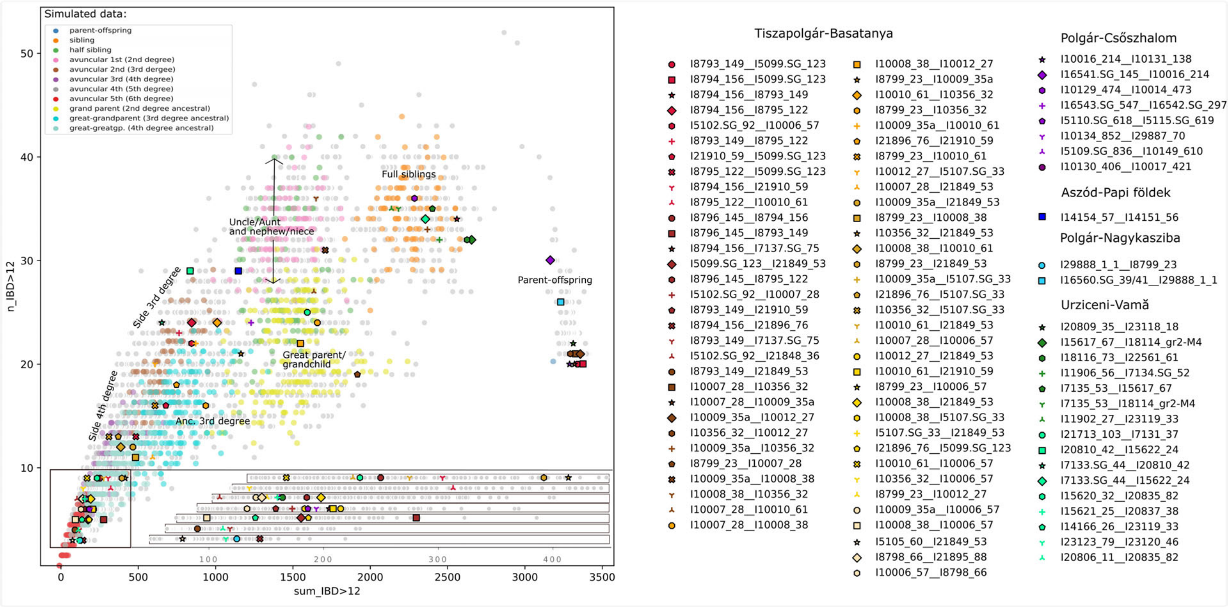

**Supplementary Figure 16. IBD scatterplot showing pairs of individuals with over 100 cM segment sharing.**

Gray dots in the background indicate actual published data pairs in the dataset. Genetic relationship was estimated through the simulated data of Ringbauer et al.[48], shown in colors, indicating type and level of kinship. Pairs of relations are presented in Supplementary Data 11.

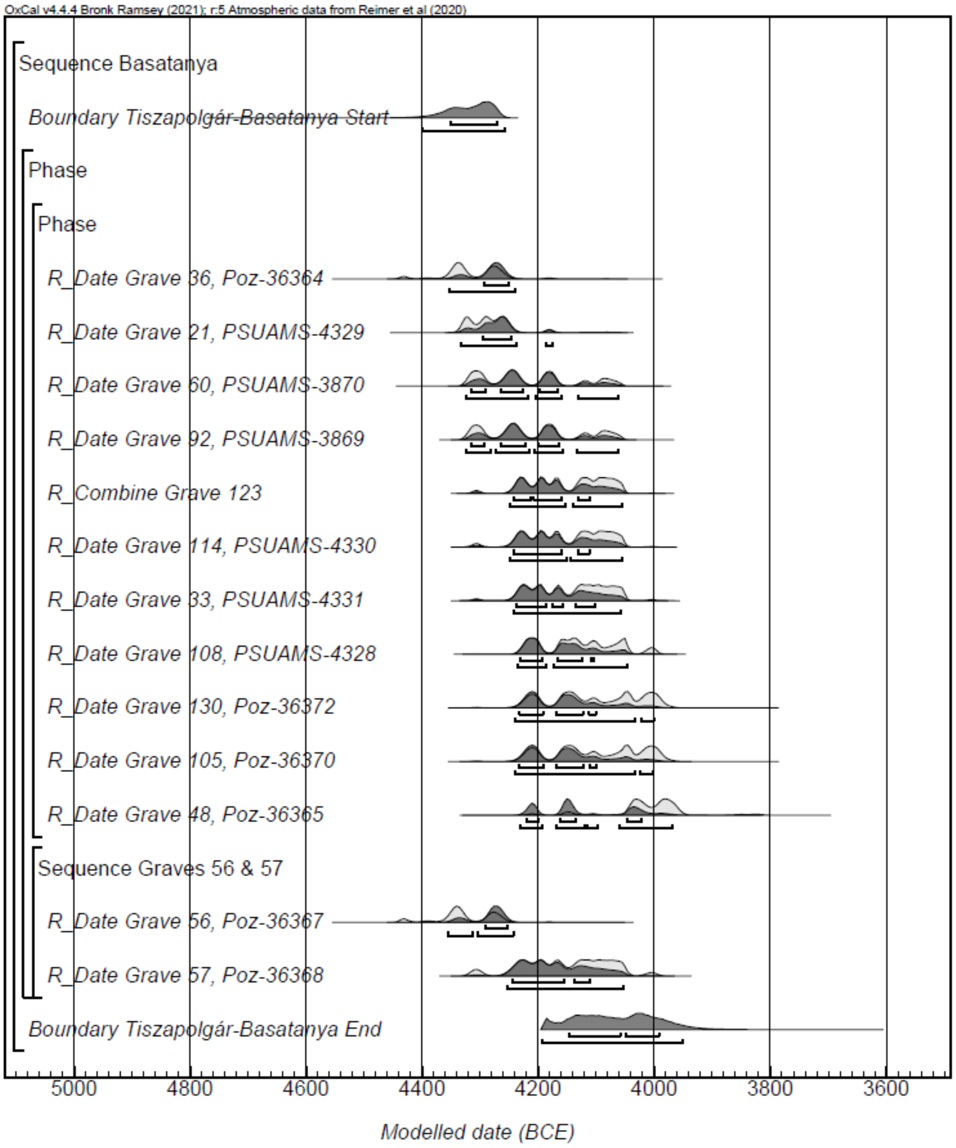

**Supplementary Figure 17. Probability distributions of radiocarbon dates from Tiszapolgár-Basatanya, Early Copper Age cemetery.**

The square brackets on the left side along with the OxCal keywords exactly define the model. The code of the OxCal modeling can be found at GitHub (github.com/ArchGenIn/Szecsenyi-Nagy_2025) and at Zenodo (DOI: 10.5281 /zenodo.15221967).

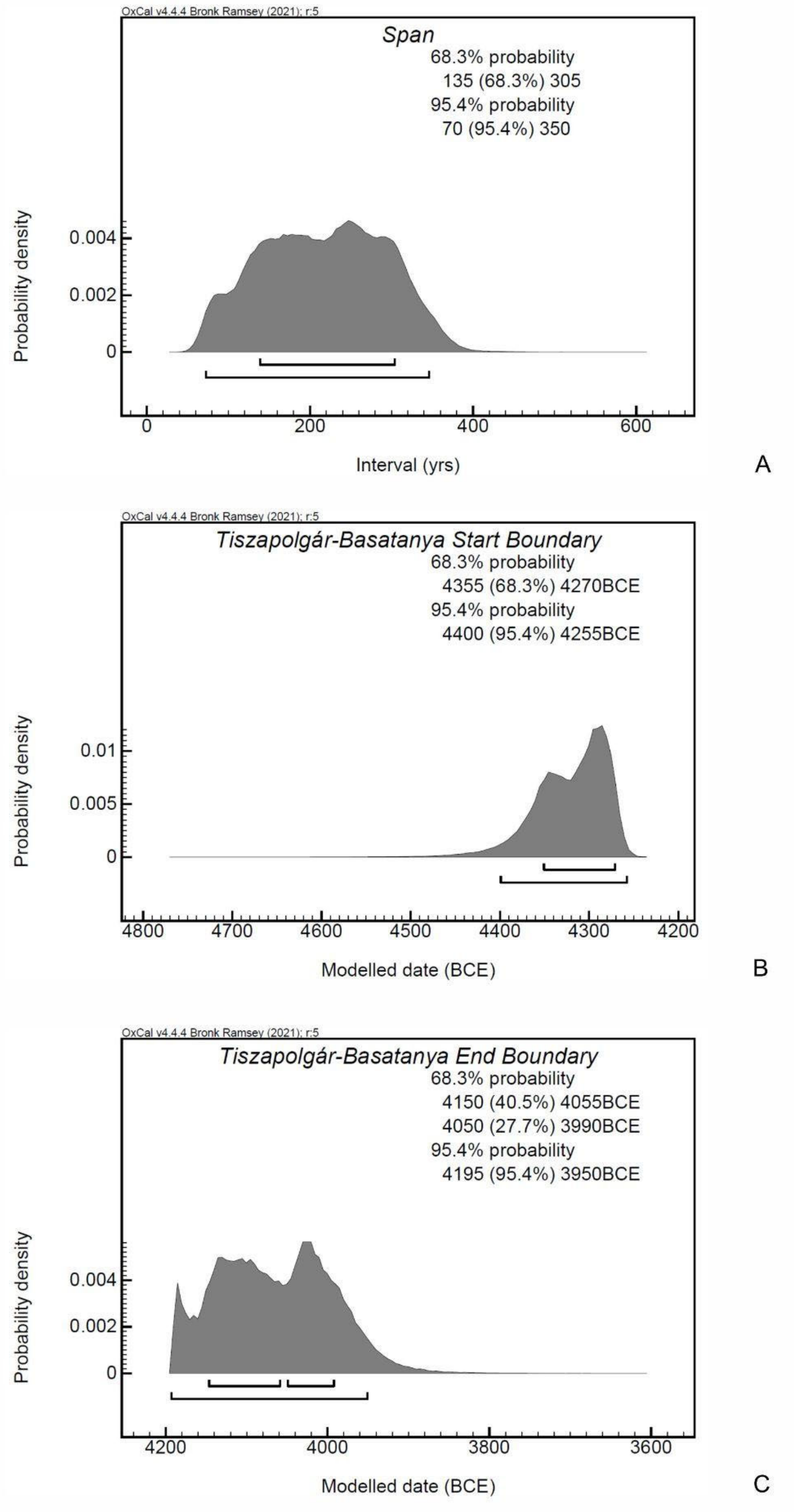

**Supplementary Figure 18. Probability distribution for the span of use (A) and the estimated start (B) and end (C) boundaries of the Tiszapolgár-Basatanya, Early Copper Age cemetery.** The code of the OxCal modeling can be found at GitHub (github.com/ArchGenIn/Szecsenyi-Nagy_2025) and at Zenodo (DOI: 10.5281 /zenodo.15221967).

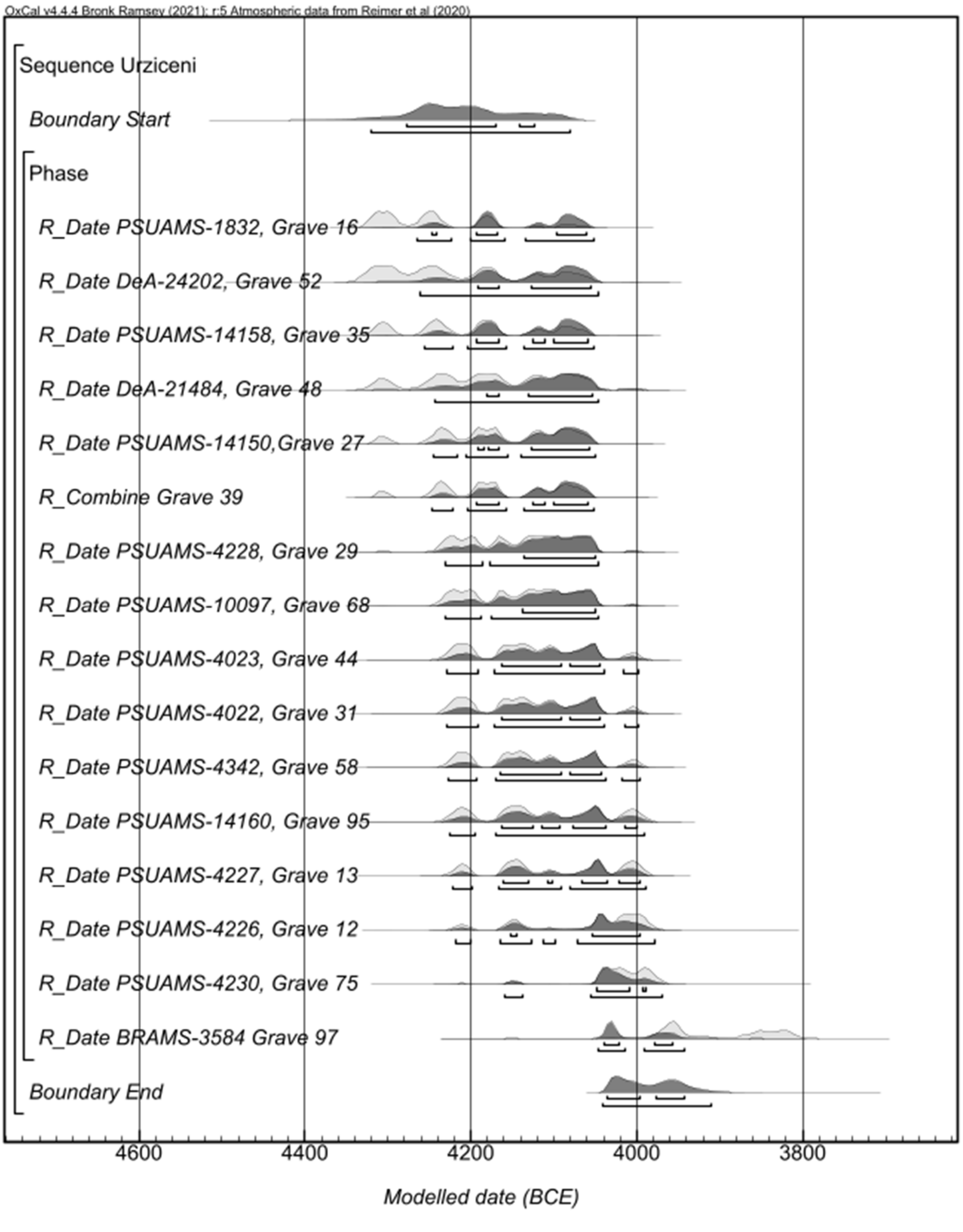

**Supplementary Figure 19. Probability distributions of radiocarbon dates from Urziceni-Vamă, Early Copper Age cemetery.**

The square brackets on the left side along with the OxCal keywords exactly define the model. The code of the OxCal modeling can be found at GitHub (github.com/ArchGenIn/Szecsenyi-Nagy_2025) and at Zenodo (DOI: 10.5281 /zenodo.15221967).

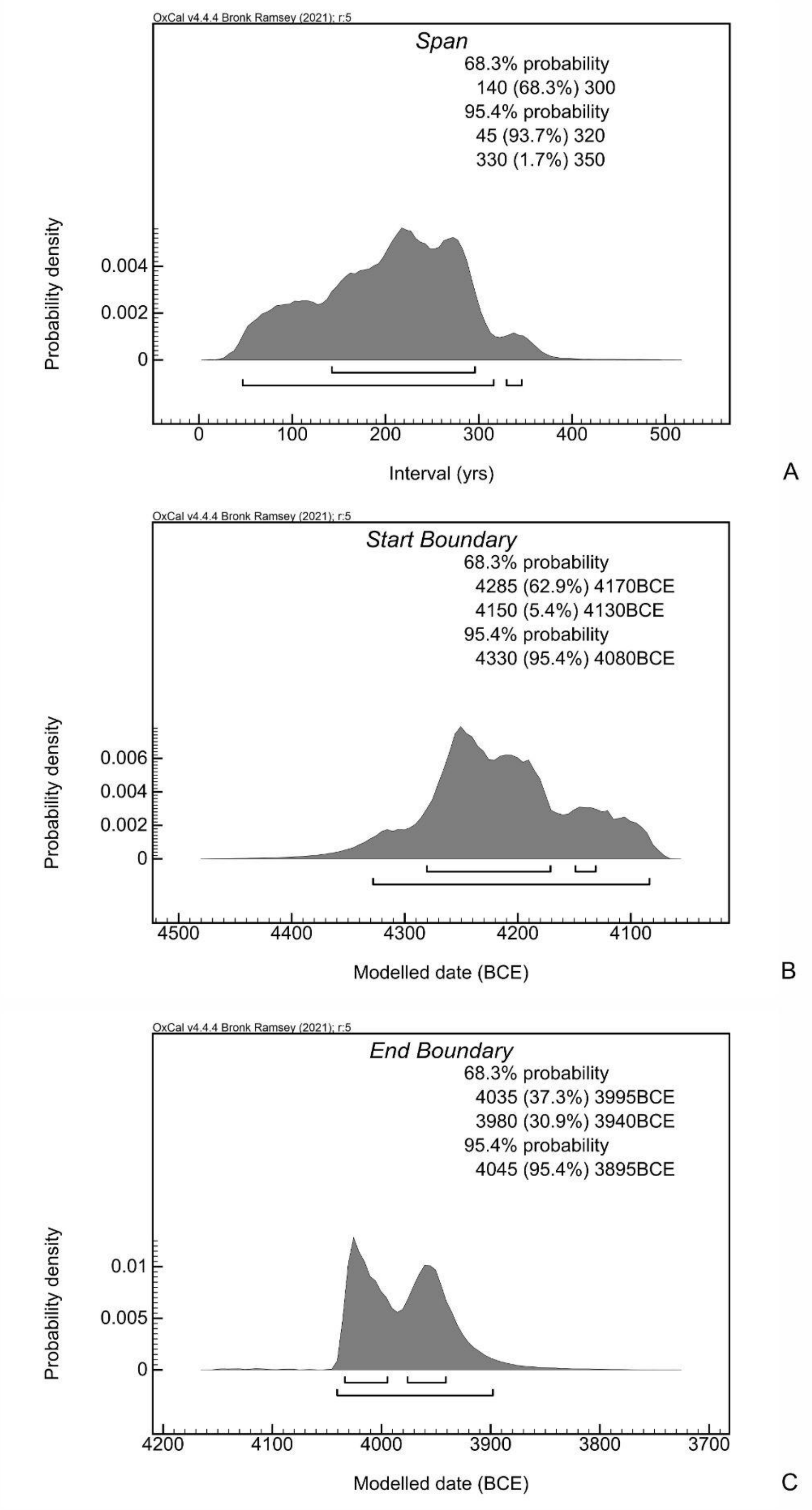

**Supplementary Figure 20. Probability distribution for the span of use (A) and the estimated start (B) and end (C) boundaries of the Urziceni-Vama.** The code of the OxCal modeling can be found at GitHub (github.com/ArchGenIn/Szecsenyi-Nagy_2025) and at Zenodo (DOI: 10.5281 /zenodo.15221967).

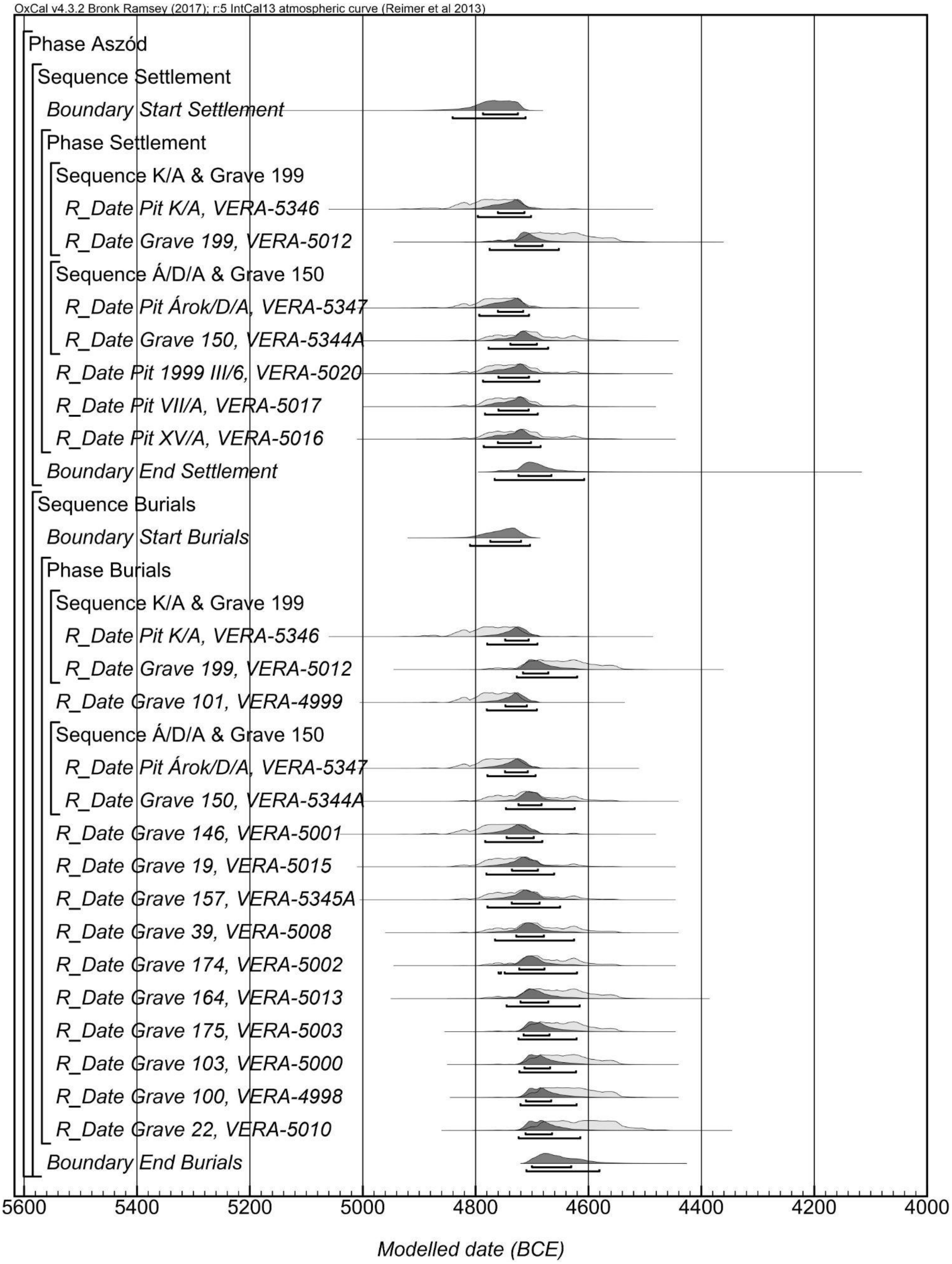

**Supplementary Figure 21. Probability distributions of radiocarbon dates from Aszód-Papi földek, Late Neolithic site.**

The square brackets on the left side along with the OxCal keywords exactly define the model. The code of the OxCal modeling can be found at GitHub (github.com/ArchGenIn/Szecsenyi-Nagy_2025) and at Zenodo (DOI: 10.5281 /zenodo.15221967).

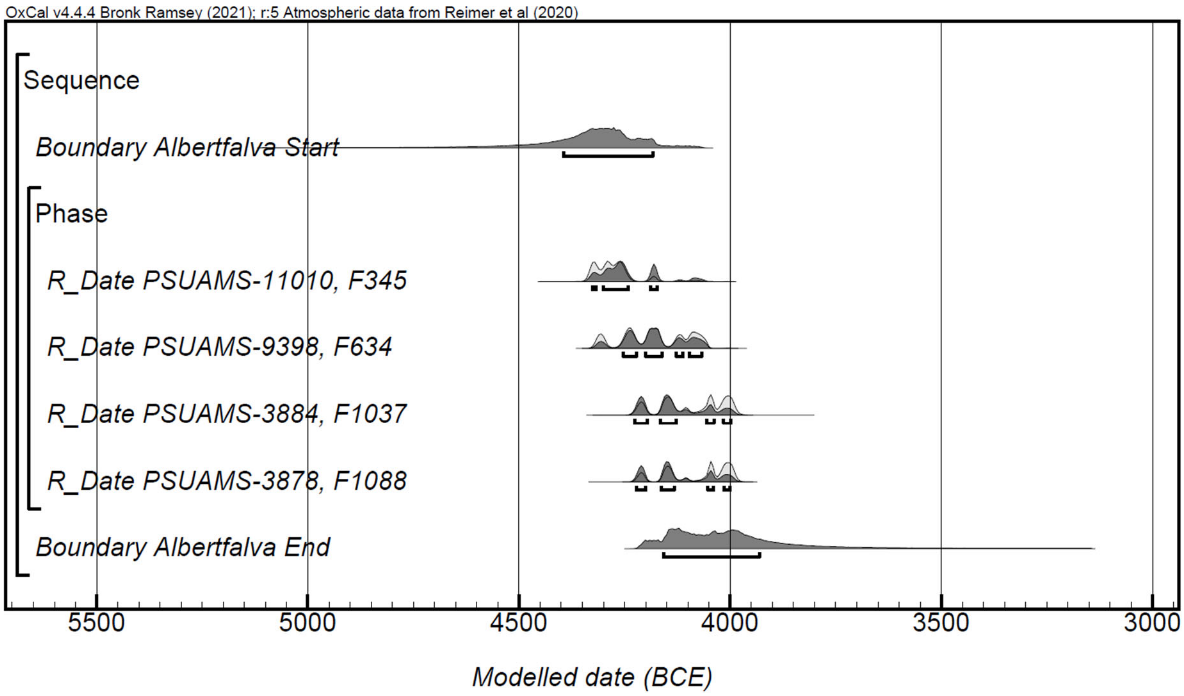

**Supplementary Figure 22. Probability distributions of radiocarbon dates from Budapest-Albertfalva-Hunyadi János út, Early Copper Age burials.**

The square brackets on the left side along with the OxCal keywords exactly define the model. The code of the OxCal modeling can be found at GitHub (github.com/ArchGenIn/Szecsenyi-Nagy_2025) and at Zenodo (DOI: 10.5281 /zenodo.15221967).

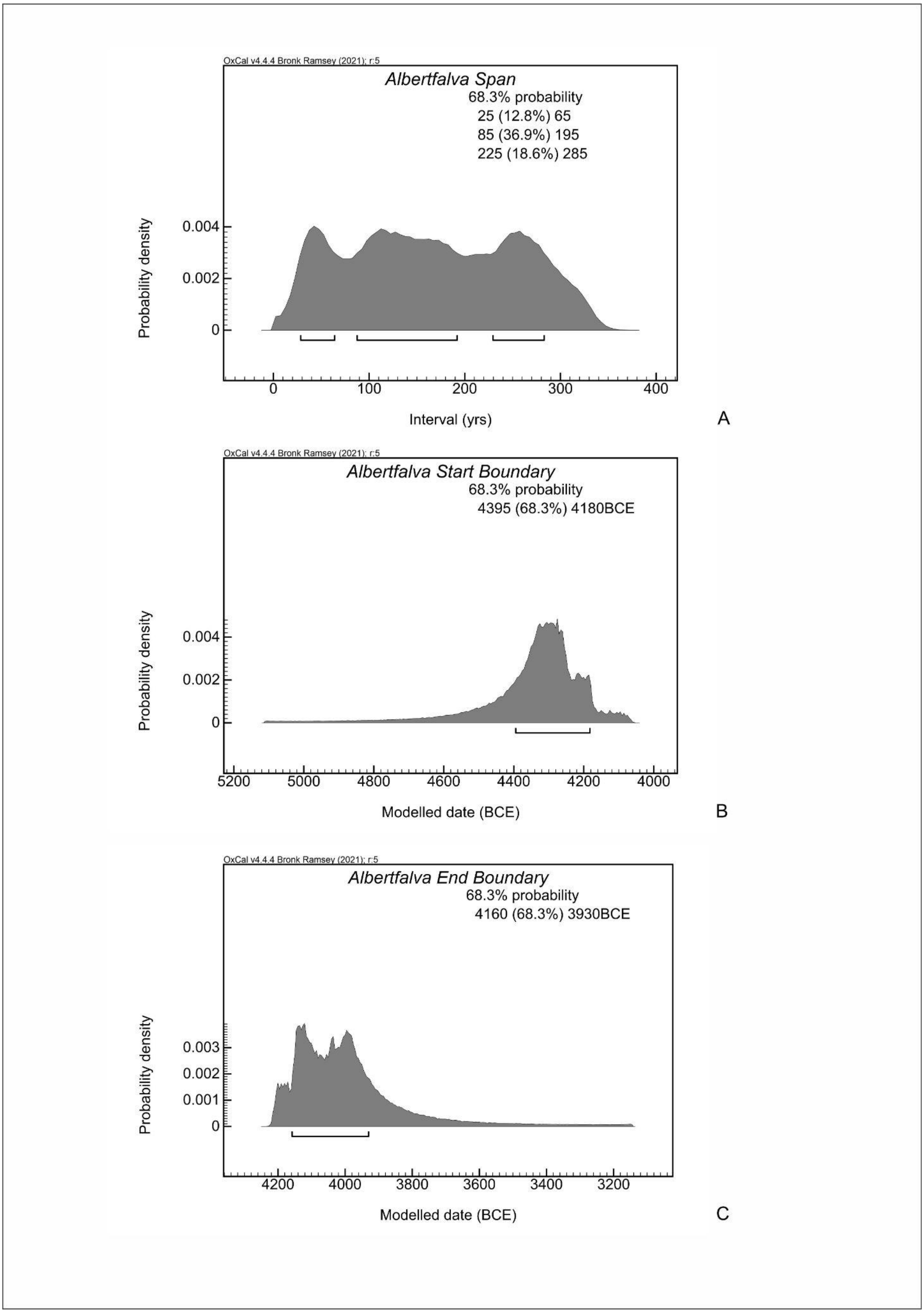

**Supplementary Figure 23. Probability distribution for the span of use (A) and the estimated start (B) and end (C) boundaries of the Budapest-Albertfalva-Hunyadi János út, Early Copper Age**

**burials.** The code of the OxCal modeling can be found at GitHub (github.com/ArchGenIn/Szecsenyi-Nagy_2025) and at Zenodo (DOI: 10.5281 /zenodo.15221967).
